# Anesthetic-Binding Induced Motion of GABA_A_ Receptors Revealed by Coarse-Grained Molecular Dynamics Simulations

**DOI:** 10.1101/2023.03.10.530555

**Authors:** Shuya Nakata, Yoshiharu Mori, Shigenori Tanaka

## Abstract

General anesthetics are indispensable in modern medicine because they induce a reversible loss of consciousness and sensation in humans. On the other hand, their molecular mechanisms of action have not yet been elucidated. Several studies have identified the main targets of some general anesthetics. The structures of γ-aminobutyric acid A (GABA_A_) receptors with the intravenous anesthetics such as propofol and etomidate have recently been determined. Although these anesthetic-binding structures provide essential insights into the mechanism of action of anesthetics, the detailed molecular mechanism of how the anesthetic binding affects the Cl^−^ permeability of GABA_A_ receptors remains to be elucidated. In this study, we performed coarse-grained molecular dynamics simulations for GABA_A_ receptors and analyzed the resulting simulation trajectories to investigate the effects of anesthetic binding on the motion of GABA_A_ receptors. The results showed large structural fluctuations in GABA_A_ receptors, correlations of motion between the amino-acid residues, large amplitude motion, and autocorrelated slow motion, which were obtained by advanced statistical analyses. In addition, comparison of the resulting trajectories in the presence or absence of the anesthetic molecules revealed a characteristic pore motion related to the gate-opening motion of GABA_A_ receptors.

## Introduction

Understanding the mechanisms of action of general anesthetics at the molecular level is important [1–6]. It provides a basis for improving existing general anesthetics and developing new ones. Despite many years of research, the mechanisms of action of general anesthetics remain unclear. Recent studies have revealed that the specifically involved molecules of some general anesthetics and some related ion channels, such as γ-aminobutyric acid A receptors (GABA_A_ receptors or GABA_A_Rs), glycine receptors, and nicotinic acetylcholine receptors, are now considered to be the primary targets of general anesthetics [5].

GABA_A_ receptors belong to a superfamily of the pentameric ligand-gated ion channels (pLGICs), which consist of five subunits (Fig. 1A) [7]. pLGICs have three domains, an extracellular domain (ECD), which forms the binding site for neurotransmitters, a transmembrane domain (TMD), which forms a small pore through which ions permeate, and an intracellular domain (ICD). pLGICs include glycine receptors, nicotinic acetylcholine receptors, the *Gloeobacter violaceus* ligand-gated ion channel (GLIC), and others, which share similar overall structures [8].

**Figure 1.**
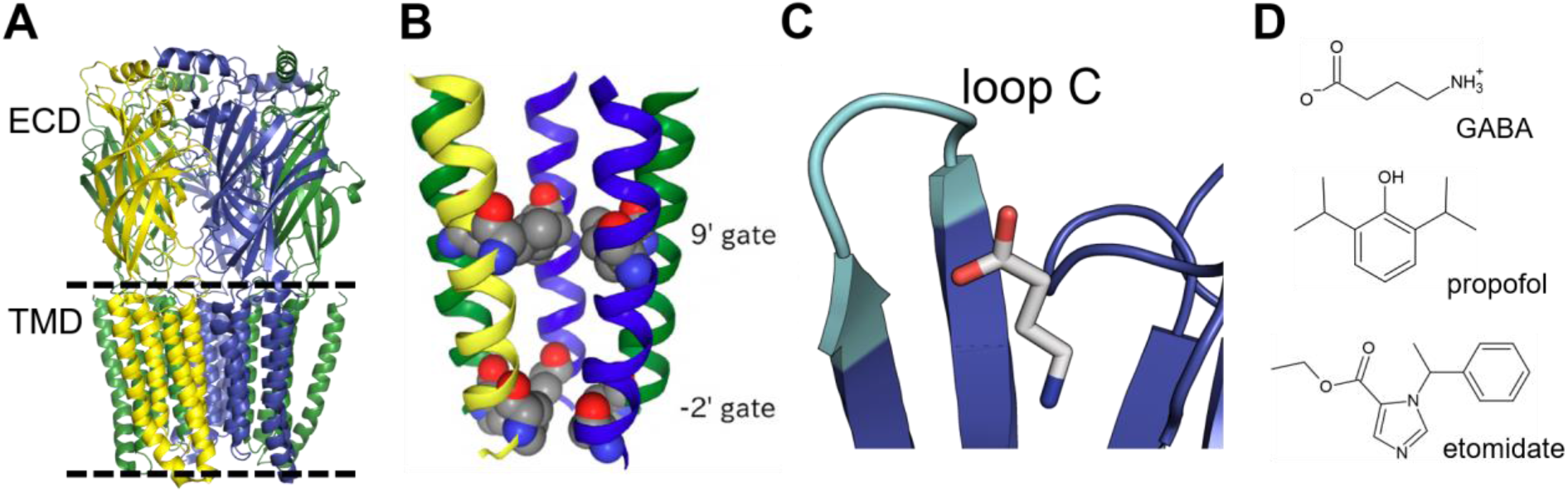
Structure of a GABA_A_R and its ligands. (A) GABA_A_Rs consist of three domains; extracellular domain (ECD), transmembrane domain (TMD), and intracellular domain (ICD, not shown). The GABA_A_R has five subunits, where the α, β, and γ subunits are colored green, blue, and yellow, respectively. (B) The pore of GABA_A_Rs has two gates, 9’ gate and −2’ gate. (C) Loop C (cyan) is located near the ligand binding site. (D) The chemical structures of GABA, propofol, and etomidate are shown.

GABA_A_ receptors are ion channels that hyperpolarize cell membranes and suppress the generation of action potentials by opening a gate in response to the binding of the neurotransmitter GABA, allowing Cl^−^ to selectively enter the cell along an electrochemical gradient. The GABA_A_R is widely distributed in the central nervous system and is the most abundant inhibitory ligand-gated ion channel in the mammalian brain [2]. There are four helices (M1-M4) in the TMD of each GABA_A_R subunit. The TMD has a small pore in the center of each of the five subunits, and the M2 helix along the pore contains two gate residues that prevent Cl^−^ permeation (Fig. 1B). The gate residues of GABA_A_Rs are a leucine residue at the 9’ position and a proline or alanine residue at the −2’ position. The 9’ gate is also called the resting gate and the −2’ gate is called the desensitization gate [9]. The subunit configuration of GABA_A_Rs in this study is one α, two β, and two γ subunits. GABA binds to the β-α interface of the ECD of GABA_A_Rs [10–13] and interacts with loop C of the β subunit (Fig. 1C). There are several α helices and β strands in the ECD of each GABA_A_R subunit. We refer to these secondary structures as α1 and β1, and so on.

GABA_A_ receptors are considered to be the central target receptors for anesthetics such as propofol and etomidate (Fig. 1D) [2]. The three-dimensional structures of complexes of anesthetics, such as propofol and etomidate, bound to GABA_A_Rs have been determined by cryogenic electron microscopy (cryo-EM) [13]. Such experimentally determined structural models have provided important insights into the molecular mechanisms of general anesthetic action [7]. In the structure of GABA_A_Rs with GABA and propofol (PDB: 6×3t) and that with GABA and etomidate (PDB: 6×3v), both propofol and etomidate bind to the same binding site at the β-α interface of the TMD [13]. However, the details of the molecular mechanism of channel gating by ligand binding, which causes Cl^−^ permeation, are still unknown.

In this study, coarse-grained molecular dynamics (MD) simulations were used to investigate the effects of ligand molecules, including general anesthetics, on the motion of GABA_A_ receptors. Dynamic properties such as the motion of each amino acid residue of the receptor, its correlation, and the motion related to channel function were investigated. We also compared the motion of GABA_A_ receptors due to different ligand molecules. These results will clarify how the binding of ligand molecules affects the function of GABA_A_ receptors.

## Computational Details

### Coarse-grained modeling of GABA_A_ receptors and ligands

We performed coarse-grained MD simulations of GABA_A_ receptors in complex with different ligands, namely GABA, propofol, and etomidate. The PDB structure of GABA_A_R with GABA (pdb id: 6×3z), GABA and propofol (pdb id: 6×3t), or GABA and etomidate (pdb id: 6×3v) [13] was used as the complex structure of the receptor and the ligands. The initial structure of each ligand-receptor complex was obtained from the OPM database [14].

A MARTINI model was used as a coarse-grained model, and a MARTINI 2.2 force field [15, 16] was used for GABA_A_ receptors. Because there are no coarse-grained models for GABA, propofol, or etomidate, the MARTINI coarse-grained model of the ligands was created as follows. First, the atoms constituting the ligand were mapped onto several coarse-grained beads. We then performed all-atom simulations of the ligand to determine parameters such as bond lengths and angles between the coarse-grained beads. Using these all-atom simulations as references, the parameters of the coarse-grained model of the ligands were estimated. We ran coarse-grained simulations of the ligand using the estimated parameters, and the parameter estimation was repeated to reproduce the results of the all-atom simulations. See Supporting Information for more details.

The coarse-grained model of the GABA_A_ receptor and ligands created was used to construct the simulation system. The ligand-receptor complex was placed in a biological membrane, which was realized by the insane program [17]. 1-palmitoyl-2-oleoylphosphatidylcholine (POPC), 1-palmitoyl-2-oleoylphosphatidylethanolamine (POPE), 1-palmitoyl-2-oleoylphosphatidylserine (POPS), phosphatidylinositol diphosphate (PIP2), and cholesterol (CHOL) were used as biomembrane constituents; POPC, POPE, POPS, POP2, and CHOL were used as lipid types (residue names) in the MARTINI model, respectively [15]. The lipid composition of the inner membrane was CHOL:POPC:POPE:POPS:POP2=40:20:20:9:1 and that of the outer membrane was CHOL:POPC:POPE=40:20:20. These lipid compositions were determined following Ingólfsson *et al*. [18]. Water was used as the solvent and the salt concentration was set to 150 mM. NaCl was used as the salt. These systems were placed in a hexagonal box and periodic boundary conditions were imposed. The individual simulation conditions are listed in Table 1.

**Table 1.** Simulation systems with different initial structures and ligand molecules.

| System | Model of protein-ligand complex | Ligands | Number of beads |
| --- | --- | --- | --- |
| 1 | GABA<sub>A</sub>R with GABA | GABA | 42,761 |
| 2 | GABA<sub>A</sub>R with GABA | None | 42,762 |
| 3 | GABA<sub>A</sub>R with GABA and propofol | GABA, propofol | 43,887 |
| 4 | GABA<sub>A</sub>R with GABA and propofol | GABA | 43,871 |
| 5 | GABA<sub>A</sub>R with GABA and propofol | None | 43,897 |
| 6 | GABA<sub>A</sub>R with GABA and etomidate | GABA, etomidate | 42,715 |
| 7 | GABA<sub>A</sub>R with GABA and etomidate | GABA | 42,721 |
| 8 | GABA<sub>A</sub>R with GABA and etomidate | None | 42,716 |

The interactions of the system were calculated using the MARTINI model [15, 16]. Each subunit of the GABA_A_ receptor was subjected to elastic networks using the Elnedyn method [19] to maintain the conformation within the GABA_A_ receptor subunit. Harmonic flat-bottom restraints were applied to allow the GABA molecule to reside in the binding region within the receptor [20] with a force constant of 5,000 kJ mol^−1^ nm^−2^. The restraints were imposed when the distance between the center of mass of the GABA molecule and the coarse-grained beads corresponding to the main chains of β-Tyr205, α-Phe65, and α-Thr130 was larger than 1.0 nm.

### MD simulation procedures

Minimization and MD simulations were performed using GROMACS 2020.4 [21]. Non-bonded interactions were calculated using electrostatic and Lennard-Jones interactions with a cutoff distance of 1.1 nm. The electrostatic interactions were calculated using the reaction-field method [22] with ε_r_ = 15 and ε_rf_ = 0. Structure minimization was performed on the system with the initial structure under each condition. Positional restraints were applied to the protein main-chain beads and ligand beads, and the force constant was set to 1,000 kJ mol^−1^ nm^−2^. The time step for numerical integration in the MD simulations was set to 20 fs. The LINCS algorithm was used to constrain the bond length [23, 24]. The temperature was controlled by Bussi’s velocity rescaling algorithm [25] with a temperature of 310 K and a time constant of 1.0 ps. Pressure was set to 1.0 bar. MD simulations for equilibration were performed as follows. Simulations were performed for 100 ns with positional restraints of 1,000, 200, 40, 8, and 0 kJ mol^−1^ nm^−2^ for the protein and ligand. The weak coupling method [26] was used as the pressure control method. The time constant of the pressure control was set to 5.0 ps and the compressibility to 10^−4^ bar^−1^. Then, equilibration simulations were performed for 100 ns using the Parrinello-Rahman algorithm [27] as the pressure control method. The time constant of the pressure control was set to 20.0 ps and the compressibility was set to 5.0× 10^−6^ bar^−1^. After the equilibration simulations were completed, production MD simulations were performed for 10 μs. The temperature and pressure control methods were the Bussi and Parrinello-Rahman methods, respectively, and the parameters used were the same as those used for equilibration.

### MD simulation analyses

We calculated root-mean-square fluctuations (RMSFs) for the MD trajectories in each condition. The RMSF of bead *i* is defined in terms of temporal average < >_*t*_ as

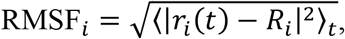

where *r_i_*(*t*) is the coordinate of bead *i* after superimposition on the reference structure at time *t* and *R_i_* is the coordinate of bead *i* in the reference structure. The average structure of the protein structures under each condition was used as the reference structure. The coordinates of the main-chain beads of the protein were used as the coordinates for the RMSF calculations.

To investigate the correlation of protein motions between each amino acid residue, the following correlation matrix *C_ij_* was calculated:

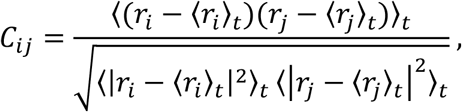

where *r_i_* and *r_j_* are the coordinates of beads *i* and *j*, respectively, at time *t*. The coordinates of the protein main-chain beads were used as the coordinates for the correlation matrix calculations.

We performed a principal component analysis (PCA) on the protein coordinates [28] for the trajectories of each simulation to identify large amplitude motions of the protein. We used the coordinates of the protein main-chain beads as the coordinates when performing the PCA.

In addition, a time-structure based independent component analysis (tICA) [29] was performed to identify slow dynamics of proteins. tICA is a method for calculating independent components (tICs) for the time series of multidimensional data such that the autocorrelation function of a tIC is maximized and uncorrelated with each other. The tICA uses the time information contained in the trajectories to extract the components of slow dynamics with high autocorrelation, which is difficult to see with PCA. For time-series data *X*(*t*), the time-delayed (Δ*t*) autocorrelation matrix 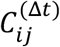 is defined as

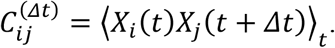

The tICs are then obtained by solving the general eigenvalue problem as in PCA. In the present analysis, Δ*t* was set to 100 ns in the present analysis.

Linear discriminant analysis with iterative procedure (LDA-ITER) [30] was also carried out to characterize the trajectories when different ligands were bound. This analysis finds the component that maximizes the ratio of the inter-ensemble variance to the intra-ensemble variance for two or more ensembles of multidimensional data. This allows for the extraction of components that characterize the differences between the ensembles.

For GABA_A_ receptors, the degree of channel gate opening is important because it affects Cl^−^ permeation. To evaluate the degree of gate opening quantitatively, we calculated the distance between non-adjacent subunits that form the channel gate. Here, we used the distance between the side-chain beads of the amino acid residues forming the 9’ and −2’ gates as the distance between non-adjacent subunits. We calculated the distances for the following non-adjacent subunit pairs: β_A_-β_C_, β_A_-α_D_, α_B_-α_D_, α_B_-γ_E_, and β_C_-γ_E_, where the subscripts correspond to the chain IDs in the PDB data [13]. Because the −2’ alanine residue of the β subunit has no side chain beads in the MARTINI model, the main-chain bead was used in the distance calculations.

## Results and Discussion

### Dynamic properties of GABA_A_ receptors

We calculated the RMSFs of the amino-acid residues of the GABA_A_ receptor from the MD simulation trajectories obtained for each condition (Fig. 2). In all cases, large RMSF values were observed at the N- and C-termini of the subunits and in some loop regions. On the other hand, the RMSF values are small at the M2 helix of the TMD, which forms the channel pore through which ions can permeate. It is possible that the structural stability of the pore corresponds to the ion permeation function of the receptor. In addition, the structural fluctuations vary from subunit to subunit. For example, the RMSF value of loop C of the β subunit, which is involved in GABA binding, was found to be particularly large compared to other regions.

**Figure 2.**
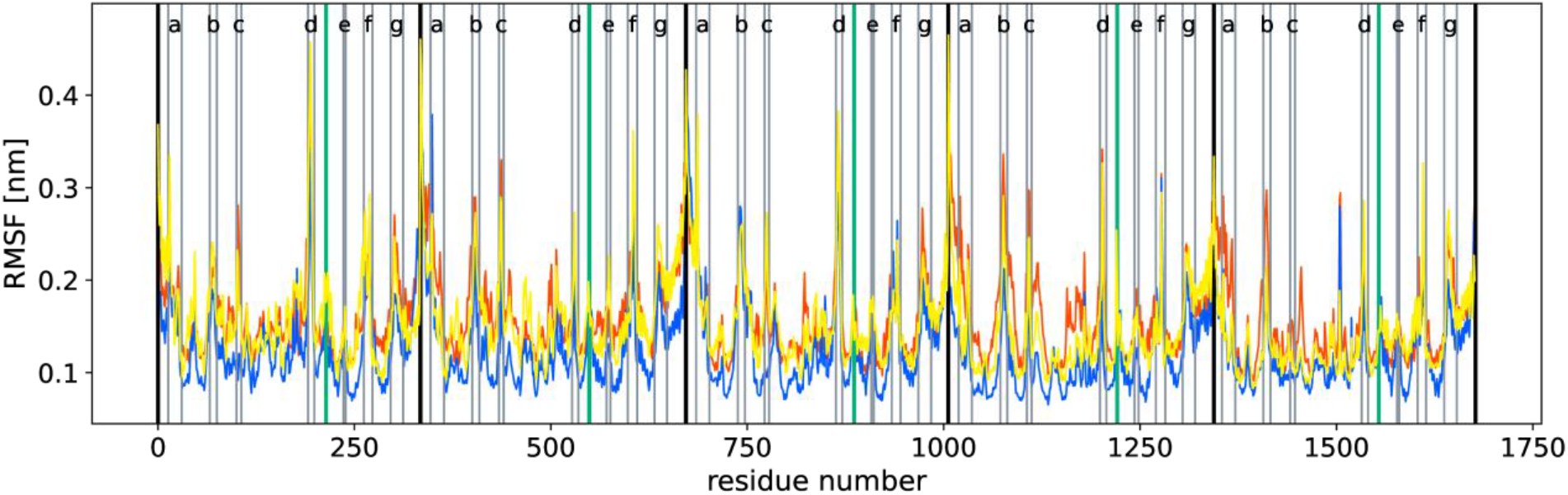
RMSF for each amino-acid residue of the GABA_A_ receptors. The red, blue, and yellow plots correspond to the RMSF of System 1, 2 (GABA), System 3-5 (GABA and propofol), and System 6-8 (GABA and etomidate), respectively. Each subunit is separated by the black vertical lines. The green vertical lines represent the boundary between the ECD and TMD. The gray lines (a-g in each subunit) indicate loop regions with large RMSF values (a: α1-β1 loop, b: β2-β3 loop, c: β4-β5 loop, d: loop C, e: M1-M2 loop, f: M2-M3 loop, and g: M3-M4 loop).

To clarify the correlation between the motion of each residue of the GABA_A_ receptor, we calculated a correlation matrix from the MD simulation trajectories for each condition (Fig. 3). The largest correlations are those between amino acid residues that form an intra-subunit. The next most notable correlations are positive correlations across domains between the M2-M3 loop of the TMD and the β 1-β2 and C loops of the ECD. These correlations suggest that these regions play an important role in the ECD-TMD communication of GABA_A_ receptors.

**Figure 3.**
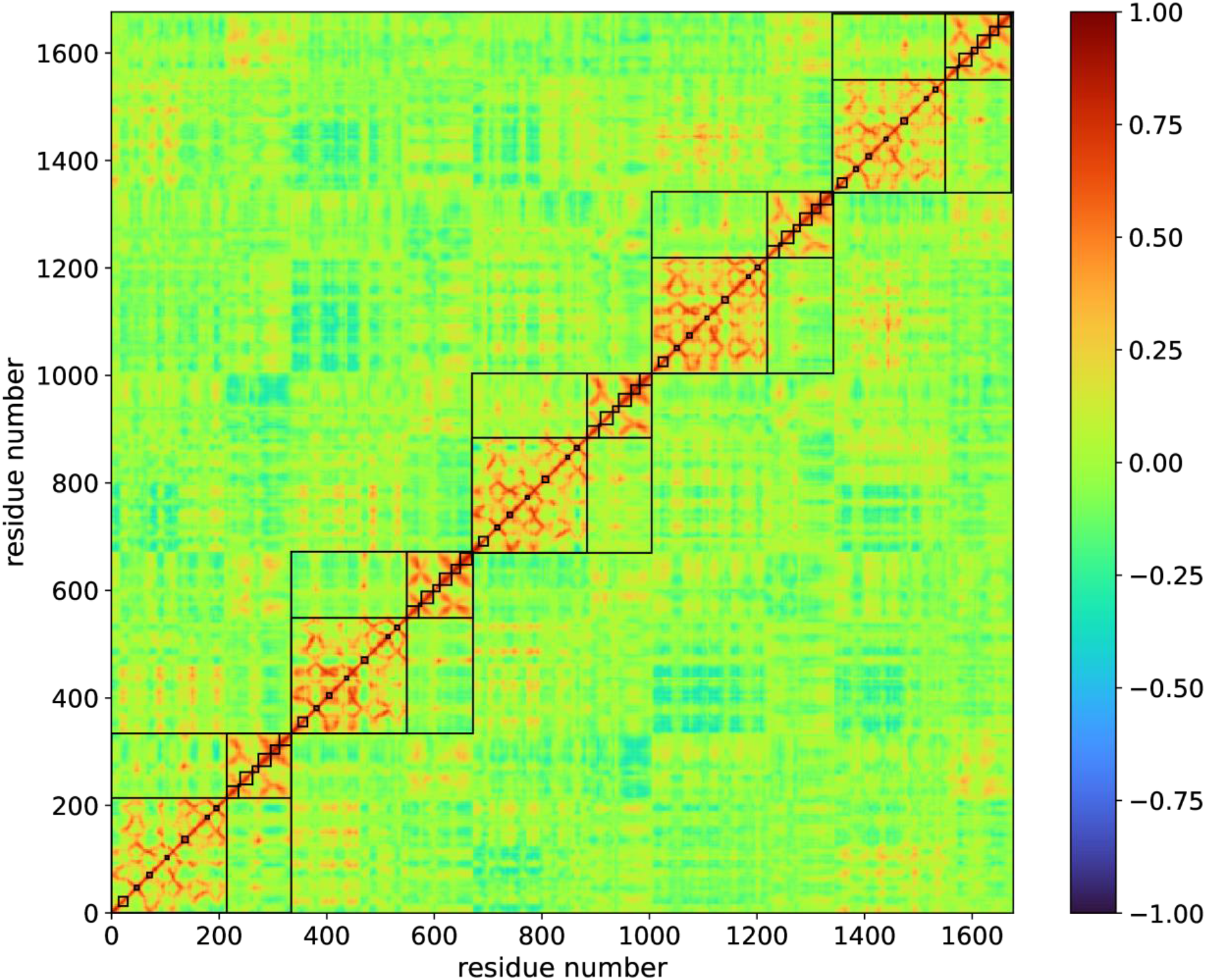
Correlation matrix for all the MD trajectories (System 1-8) of the GABA_A_ receptors. The five largest rectangles represent each subunit. Within each subunit, the lower left and upper right rectangles represent the ECD and TMD, respectively. The smaller rectangles in the lower left rectangle correspond to the α1-β1 loop, β 1-β2 loop, β2-β3 loop, β4-β5 loop, Cys-loop, β8-β9 loop, and C loop from bottom left to top right. The smaller ones in the upper right rectangle correspond to the M1 helix, M1-M2 loop, M2 helix, M2-M3 loop, M3 helix, M3-M4 loop, and M4 helix from bottom left to top right.

The PCA was performed by using the trajectories of the MD simulations under each condition. The displacements corresponding to the top three principal components and the projection of the trajectories onto these principal components for the system of GABA_A_Rs with GABA and propofol are shown in Figs. 4 and 5 (for the other systems, see Figs. S12-S15). In PC1-PC3, most of the components were found to be rotational motions of subunits or domain units. The main components of PC1 represent the rotations of the ECD of the β subunit and the ECD of the α subunit. Their rotations are in opposite directions to each other. In PC2 and PC3, we observed the rotation of the upper ECD of each subunit. In addition, the rotations of the M4 helix and the M2-M3 loop in the TMD were also observed.

**Figure 4.**
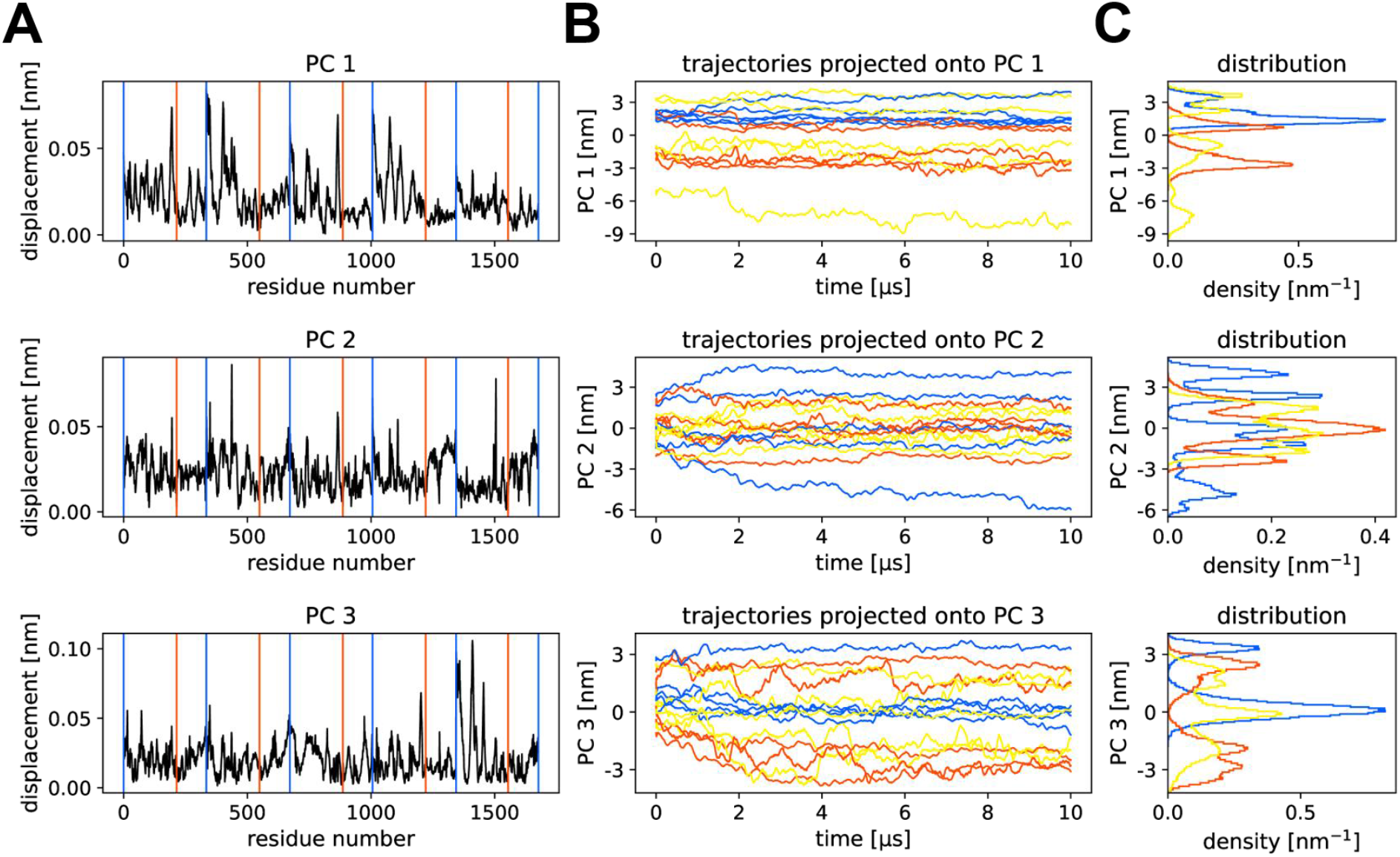
Results of the PCA for System 3-5 (GABA_A_R with GABA and propofol). (A) The displacement of each residue is plotted for the first three PCs (The explained variance ratio of PC1, PC2, and PC3 is 0.167, 0.086, and 0.077, respectively). The blue and red vertical lines represent the subunit and ECD-TMD boundary, respectively. (B, C) The trajectories projected onto each PC and their distribution are shown. The five trajectories obtained from System 3 (yellow, GABA_A_R with GABA and propofol), System 4 (red, GABA_A_R with GABA), and System 5 (blue, GABA_A_R without ligands) were projected onto the PCs calculated by the trajectories of System 3-5.

**Figure 5.**
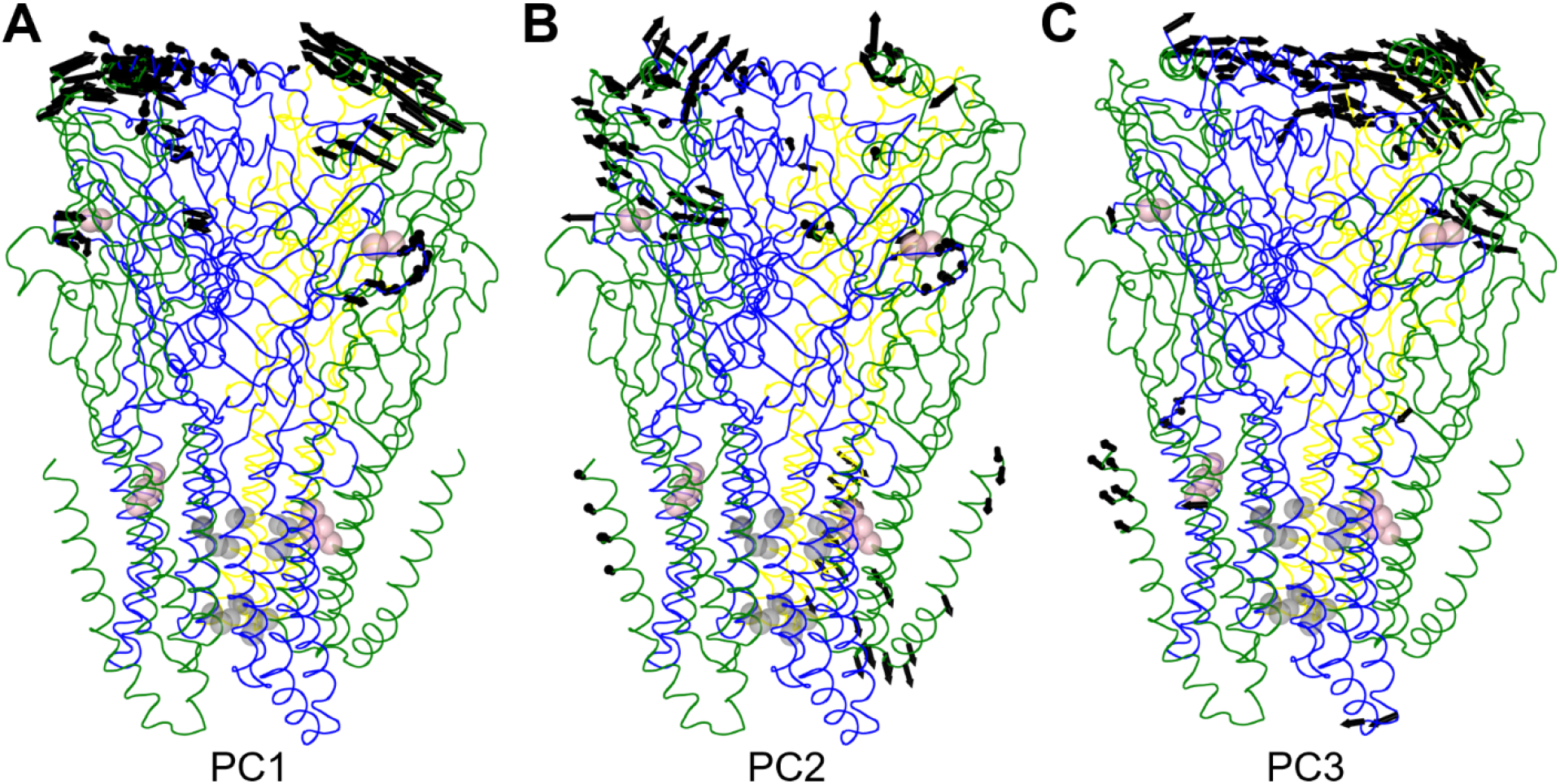
(A) PC1, (B) PC2, and (C) PC3 for GABA_A_Rs with GABA and propofol. The largest 100 components of each PC are shown by black arrows. The gray and pink spheres show the gate residues of the pore and the ligands, respectively. The α, β, and γ subunits are colored green, blue, and yellow, respectively.

We also carried out tICA for the trajectories from the MD simulations for each condition. The structural displacements and the projection of the trajectories onto the top three tICs for all the trajectories are shown in Figs. 6 and 7 (for the tICA of System 1, 2, System 3-5, and System 6-8, see Figs. S16-S21). tIC1-tIC3 show little rotation for loop C, the top of the ECD surface, or the M4 helix while such rotation was observed in the PCA. On the other hand, the tICs contain components corresponding to the M2 helix, the M2-M3 loop, the ECD-TMD interface, and the ECD subunit interface. tIC1 and tIC2 also contain large components corresponding to the motion of the M2 helix, which includes the leucine residue at the 9’ gate of the β subunit. Furthermore, focusing on the motion around the 9’ gate (β-L253), the β-L253 residue moves in the opposite direction to β-L259 in tIC1 and tIC2. This movement corresponds to the tilting of the M2 helix. Therefore, the component of slow dynamics is considered to be a characteristic motion related to the channel function of GABA_A_ receptors.

**Figure 6.**
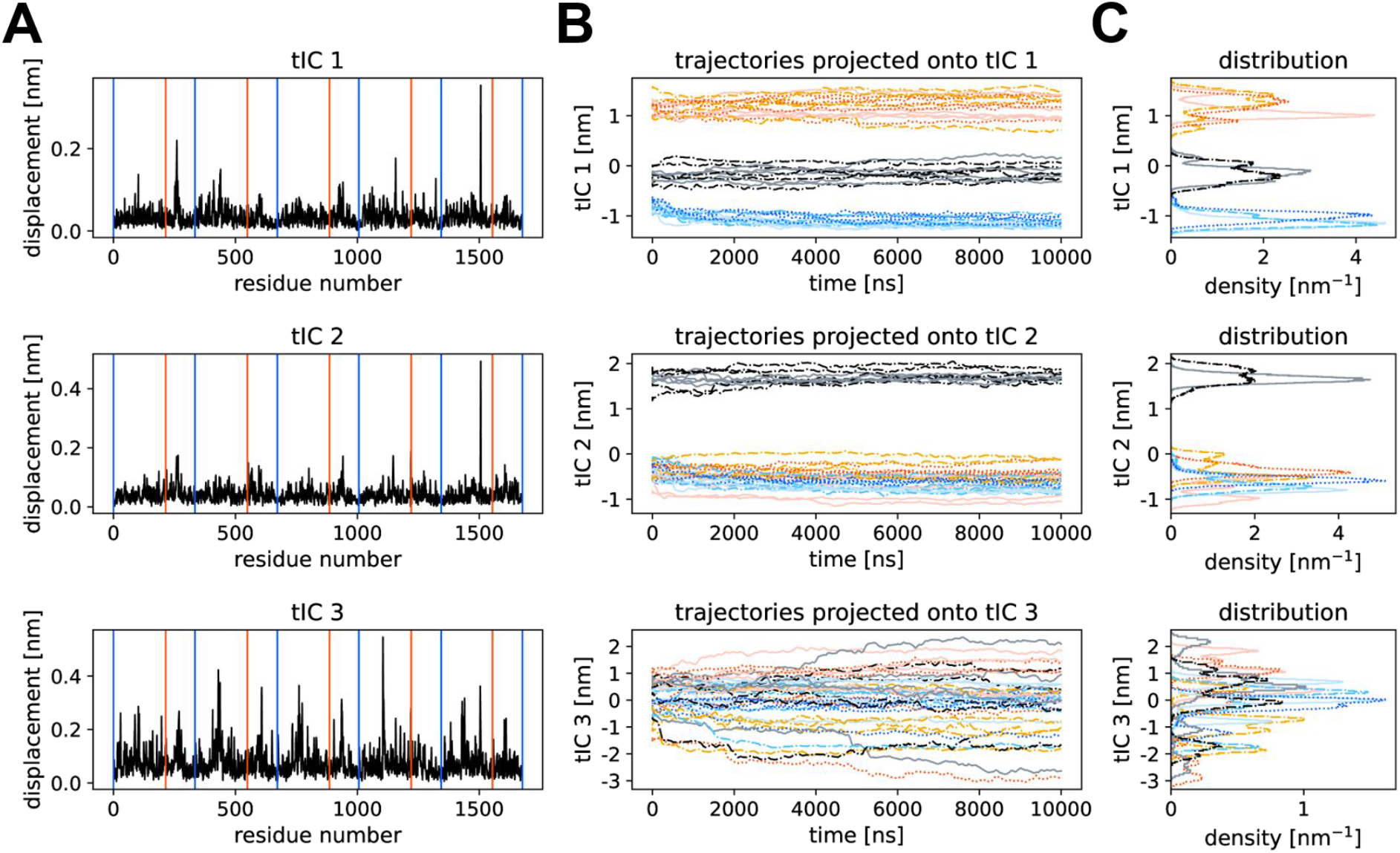
Results of the tICA obtained from all the trajectories. (A) The displacement of each residue is plotted for the first three tICs. The blue and red vertical lines represent the subunit and ECD-TMD boundaries, respectively. (B, C) The trajectories projected onto each tIC and their distribution are shown. The five trajectories obtained from System 1 (dash-dotted black, GABA_A_R with GABA), System 2 (solid gray, GABA_A_R without GABA), System 3 (dotted red, GABA_A_R with GABA and propofol), System 4 (dash-dotted orange, GABA_A_R with GABA), System 5 (solid light pink, GABA_A_R without ligands), System 6 (dotted blue, GABA_A_R with GABA and etomidate), System 7 (dash-dotted cyan, GABA_A_R with GABA), and System 8 (solid light cyan, GABA_A_R without ligands) were projected onto the tICs calculated from all the trajectories.

**Figure 7.**
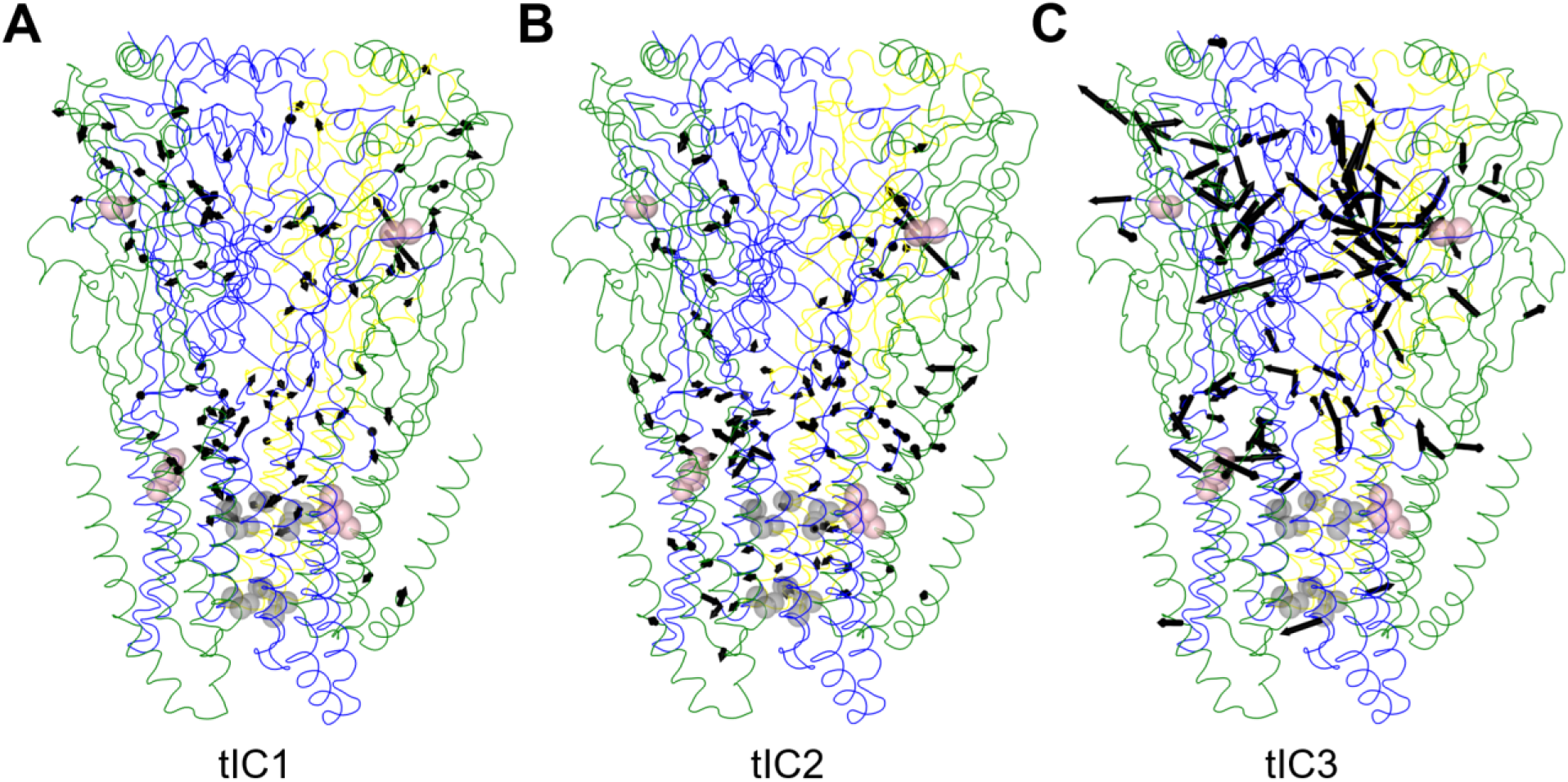
(A) tIC1, (B) tIC2, and (C) tIC3 for GABA_A_Rs. The largest 100 components of each tIC are shown by black arrows. The gray and pink spheres show the gate residues of the pore and the ligands, respectively. The α, β, and γ subunits are colored green, blue, and yellow, respectively.

### Changes in the dynamic properties of GABA_A_ receptors upon ligand binding

To characterize the motion of the GABA_A_ receptor in the presence or absence of each ligand, we performed LDA-ITER analyses on the GABA_A_ receptor (Figs. 8 and 9). We also calculated the distances between non-adjacent subunits corresponding to the channel gate for each trajectory to evaluate the degree of opening of the 9’ and −2’ gates quantitatively. Figure 10 shows the time series of the inter-subunit distances and the distribution of the distances for each ligand condition.

**Figure 8.**
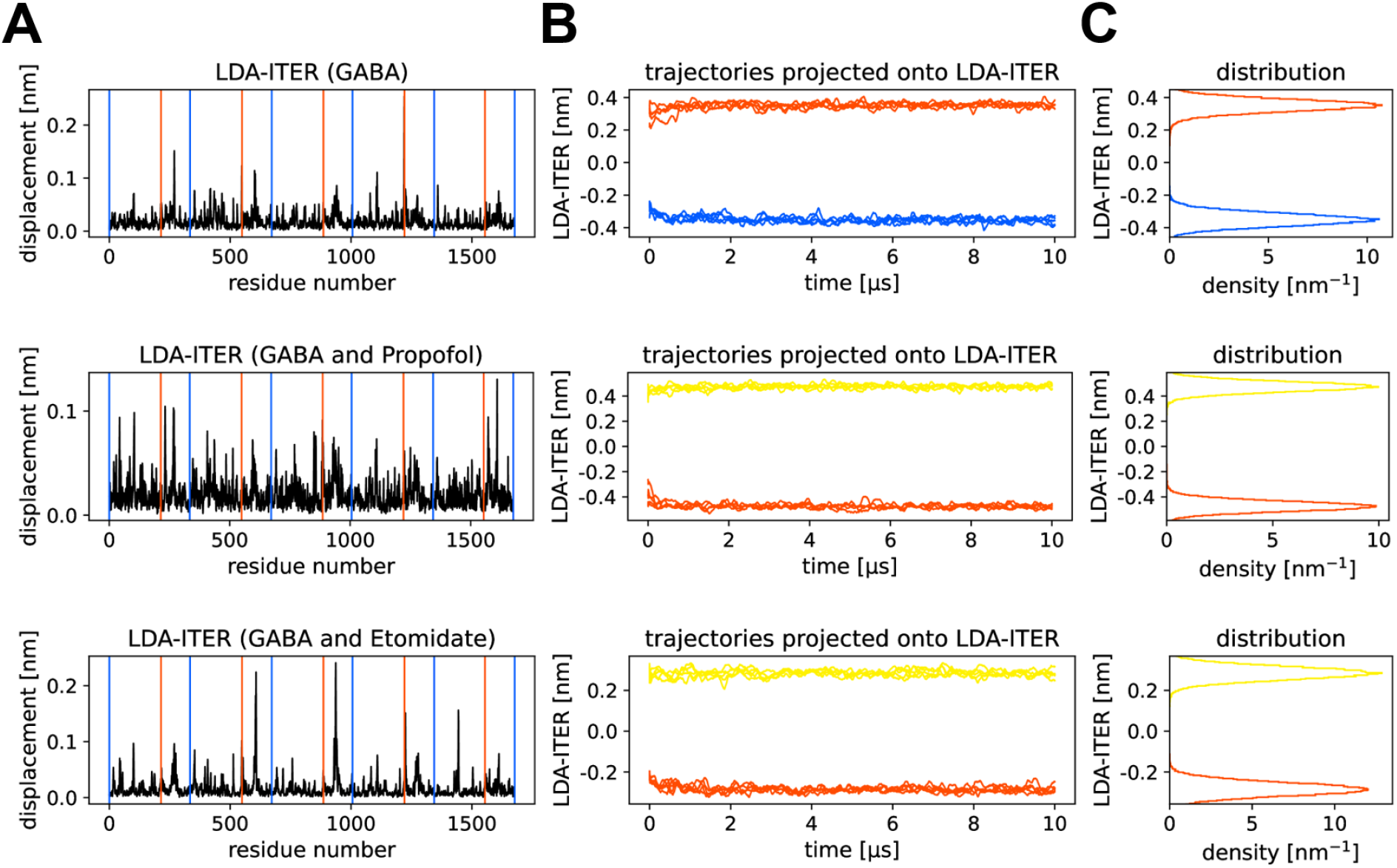
Results of the LDA-ITER analyses for System 1, 2 (GABA_A_R with GABA), System 3-5 (GABA_A_R with GABA and propofol), and System 6-8 (GABA_A_R with GABA and etomidate). (A) The displacement of each residue is plotted for the results of the LDA-ITER analyses. The blue and red vertical lines represent the subunit and ECD-TMD boundaries, respectively. (B, C) The trajectories projected onto the axis of the LDA-ITER analyses and their distribution are shown. The trajectories obtained from System 1 (red) and System 2 (blue, GABA_A_R without GABA) were projected onto the axis of the LDA-ITER results calculated from the trajectories of System 1, 2. The trajectories obtained from System 3 (yellow) and System 4 (red, GABA_A_R with GABA) were projected onto the axis of the LDA-ITER results calculated from the trajectories of System 3-5. The trajectories obtained from System 6 (yellow) and System 7 (red, GABA_A_R with GABA) were projected onto the axis of the LDA-ITER results calculated from the trajectories of System 6-8.

**Figure 9.**
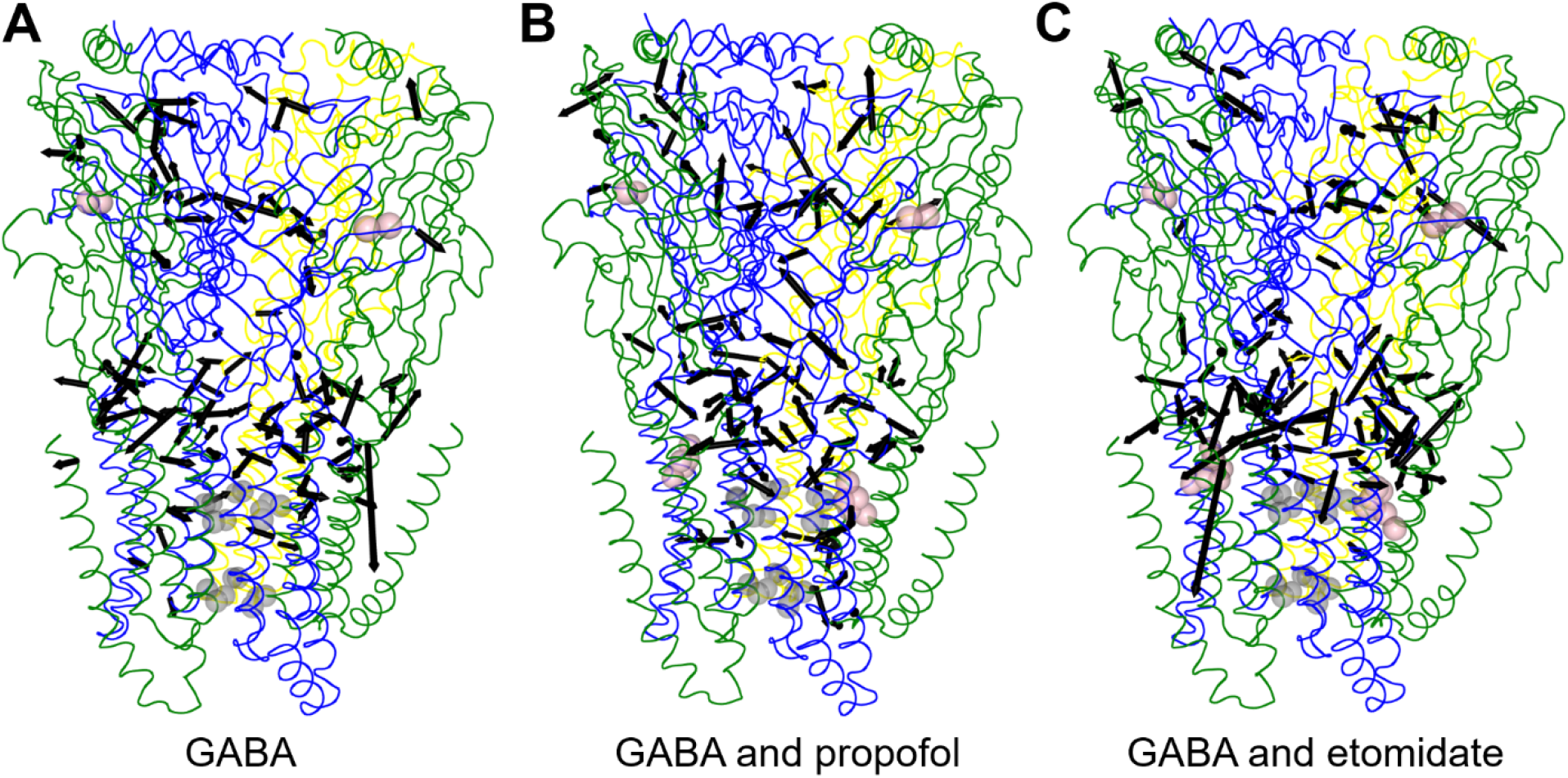
Results of the LDA-ITER analyses for (A) System 1, 2 (GABA_A_R without GABA), (B) System 3-5 (GABA_A_R with GABA), and (C) System 6-8 (GABA_A_R with GABA). The largest 100 components are shown by black arrows. The gray and pink spheres show the gate residues of the pore and the ligands, respectively. The α, β, and γ subunits are colored green, blue, and yellow, respectively.

**Figure 10.**
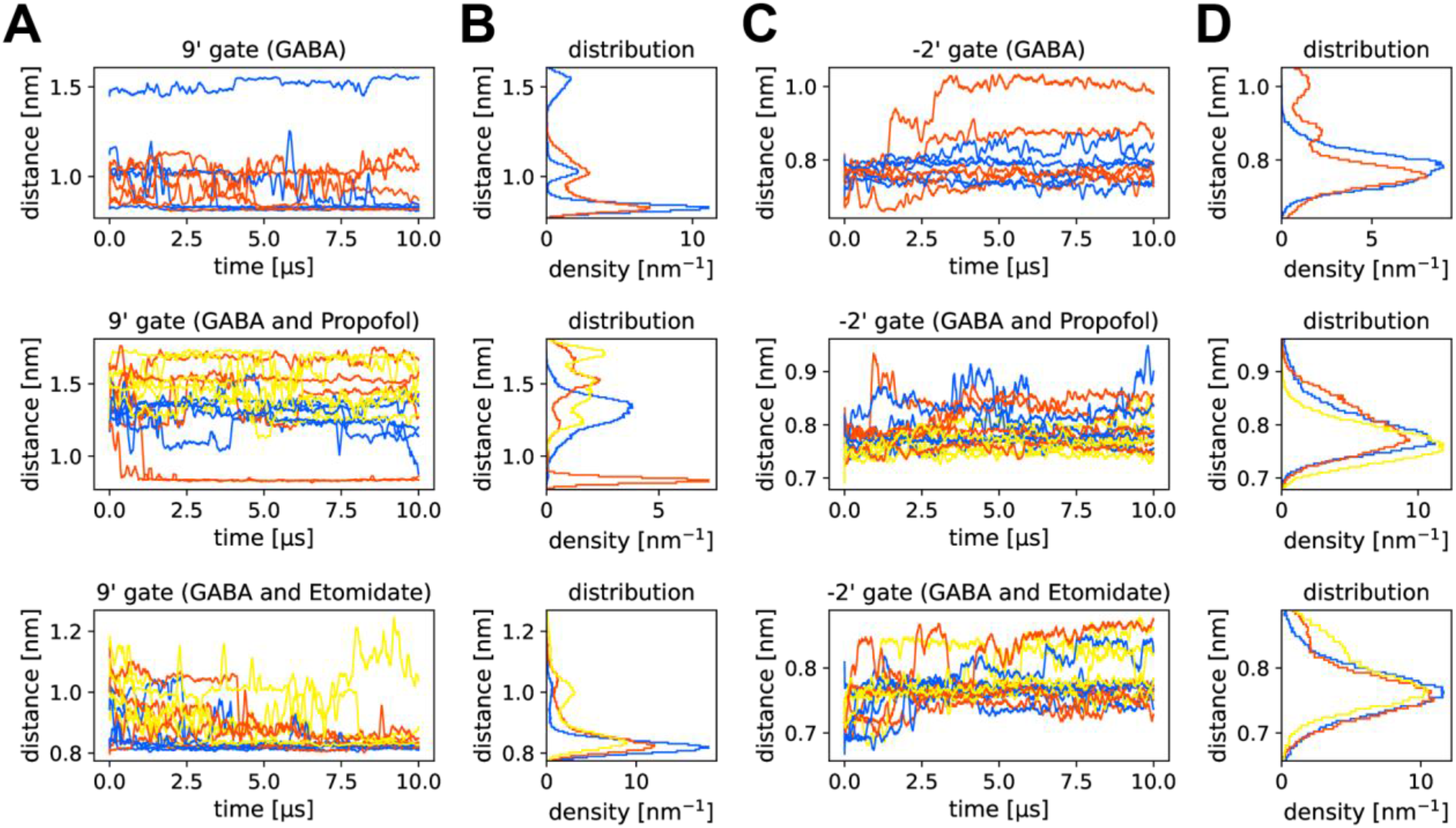
Time series and distribution of distances between the amino-acid residues of the non-adjacent subunits. The time series of the distances and the distance distribution for the 9’-gate pairs are shown in (A) and (B), and those for the −2’-gate pairs are shown in (C) and (D). The yellow, orange, and blue curves correspond to the results of System 3 and 6 (GABA_A_R with GABA and propofol or GABA and etomidate), System 1, 4, and 7 (GABA_A_R with GABA), and System 2, 5, and 8 (GABA_A_R without ligands), respectively.

The results of the LDA-ITER analysis for propofol shown in Figs. 8 and 9 indicate that the M2, M3 helices and M2-M3 loop of the β subunit and the M1, M2 helices and M2-M3 loop of the α subunit are the most important components characterizing the presence of propofol. In particular, at the propofol binding site in the β-α interface, we observed a component of movement toward the propofol binding region. This could be due to the movement of the amino-acid residues around the binding site in the direction of the gap created by the exclusion of propofol in the trajectories without propofol. The binding features of propofol were also observed at the ECD-subunit interface and the ECD-TMD interface, which are distant from the binding site. This suggests that the binding of propofol affects a large region of the subunit interfaces. Figure 10 shows the distances between the amino-acid residues of the non-adjacent subunits for the propofol-bound trajectories. The distance between the residues of the 9’ gate was maintained at about 1.5 nm in the presence of propofol. On the other hand, the distances between the residues in the trajectories without propofol tended to decrease with time. These results support the possibility that the binding of general anesthetics stabilizes the 9’ open-like conformation [13].

The results of the LDA-ITER analysis for etomidate shown in Figs. 8 and 9 indicate that, as in the case of propofol, the M2, M3 helices and M2-M3 loop of the β subunit and the M1, M2 helices and M2-M3 loop of the α subunit are important components that characterize the presence of etomidate. Movement toward the etomidate binding region was observed. The amino-acid residues around the binding site moved in the direction of the gap created by the exclusion of etomidate. The TMD component near the binding site is larger for etomidate than for propofol. This could reflect the difference in molecular size between propofol and etomidate.

## Conclusions

We performed coarse-grained MD simulations on GABA_A_ receptors, which are considered to be the main targets of anesthetics such as propofol and etomidate, to elucidate the molecular mechanism of general anesthetic action. The dynamic properties of GABA_A_ receptors were analyzed to determine the effects of ligand binding by GABA, propofol, and etomidate on GABA_A_ receptor movement.

RMSF and PCA calculations for the GABA_A_R trajectories obtained from the MD simulations allowed us to identify regions of large structural fluctuations in the receptor. In addition, the correlation matrix calculation and tICA for the MD trajectories of the receptor revealed the correlation between the motions of the amino acid residues and the components of the slow dynamics with large autocorrelation, respectively. In particular, tICA revealed the slowest motions at the subunit and ECD-TMD interfaces. Furthermore, we observed the movement of the 9’-leucine residue of the β subunit toward the channel pore, which is thought to directly affect the channel function. We found that this movement is caused by the tilting of the M2 helix of the receptor.

In addition to the PCA and tICA calculations, LDA-ITER analyses were performed to investigate the effects of ligand binding on the movement of GABA_A_ receptors. These analyses allowed us to extract the dynamic components that characterize the presence or absence of ligands. We also calculated the distances between the amino-acid residues in non-adjacent subunits corresponding to the channel gate. We identified how ligand binding regulates the degree of the gate opening. These analyses suggest that ligand binding to the receptor affects not only the vicinity of the ligand binding site, but also the entire subunit interface. All-atom molecular dynamics simulations would be useful to explain in detail the overall structural fluctuations caused by ligand binding.

Our coarse-grained MD simulations could provide information about the dynamic properties of the GABA_A_ receptor and the effects of ligand binding on its motion, which will lead to a better understanding of the GABA_A_ receptor and the mechanism of action of general anesthetics. These findings will also be useful for the improvement of general anesthetics and the development of new drugs.

## Supporting Information

Coarse-grained modeling for GABA, propofol, and etomidate; additional results for the coarse-grained molecular dynamics simulations.

## Acknowledgments

We acknowledge the Grants-in-Aid for Scientific Research from the Ministry of Education, Culture, Sports, Science and Technology (MEXT) (No. JP17H06353) and Japan Society for the Promotion of Science (JSPS) (Nos. JP18K03825, JP21K06098, JP21K06113), and MEXT Quantum Leap Flagship Program (No. JPMXS0120330644). The computation was partly performed using Research Center for Computational Science, Okazaki, Japan (Project: 20-IMS-C261, 21-IMS-C013, 22-IMS-C014).

