## Supporting Information for "Anesthetic-Binding Induced Motion of GABA_A_ Receptors Revealed by Coarse-Grained Molecular Dynamics Simulations"

### Coarse-grained modeling of GABA, propofol, and etomidate

Because there are no MARTINI force fields for GABA, propofol, or etomidate, their parameters for coarse-grained MD simulations were determined as follows. First, groups of atoms in the all-atom model were determined to be mapped onto coarse-grained beads according to the MARTINI coarse-grained model for molecules such as amino acids and lipids [1-3]. The bead type was also determined based on the chemical properties of the atoms in the coarse-grained beads.

The structures and force fields of GABA, propofol, and etomidate were obtained from the ATB database [4], which provides those of various ligand molecules. The structure and force field of octanol used as a solvent were also obtained from the database. We used the GROMOS 54A7 force field [5]. All-atom MD simulations were performed using GROMACS 2020.4 [6]. We used water as the solvent for GABA and octanol for propofol and etomidate. The MD simulations were run for 50 ns at 300 K for these systems.

The all-atom model was mapped to the coarse-grained model by using the geometric center of the atomic groups as the coordinate of the coarse-grained beads. After mapping, the parameters of the ligands were determined by using the distribution of bond distances, angles, and dihedral angles for these beads.

Coarse-grained MD simulations were performed using these parameters for a system with one ligand molecule in solvent. After performing structure optimization, a 1.0 ns equilibration run and a 50 ns production run were performed at a temperature of 300 K. The MD simulation was performed using GROMACS 2020.4, and the force field was MARTINI 2.2. The obtained distributions of bond distances, angles, dihedral angles, and solvent-accessible surface area (SASA) in the coarse-grained model were compared with those in the all-atom model. For the calculation of SASA, the Lennard-Jones parameters in the MARTINI force field were used as the radius of the beads in the coarse-grained model ($2^{1/6}\sigma$). Free energy calculations were also performed to obtain log*P* values (*P* stands for a partition coefficient). Umbrella sampling [7] was used to evaluate the free energy. The reaction coordinates for GABA were defined as the path from the water layer to the octanol layer (Fig. S1), with the center of the water layer as the origin. We used the path from the octanol layer to the water layer as the reaction coordinates for propofol and etomidate, with the center of the octanol layer as the origin.

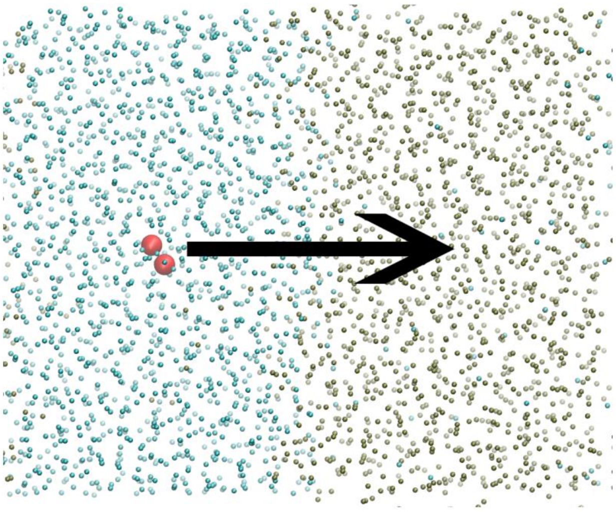

Figure S1. Reaction coordinate for umbrella sampling simulations of GABA.

The ligand properties of the resulting coarse-grained model were evaluated. If the model did not sufficiently reproduce those of the all-atom model in the evaluation, we returned to the mapping step and adjusted the parameters manually.

The results of the coarse-grained MD simulation using the obtained GABA parameters are shown in Figs. S2-S4. Although the coarse-grained model for GABA did not reproduce the bimodal distribution seen in the all-atom model, it reproduced the average distance between the beads. The coarse-grained MD simulation also reproduced the SASA distribution with sufficient accuracy. The log*P* value calculated from the free energy profile was −3.39. This value is close to the experimental value of −3.17, which indicates that the coarse-grained model reproduced the polarity of GABA.

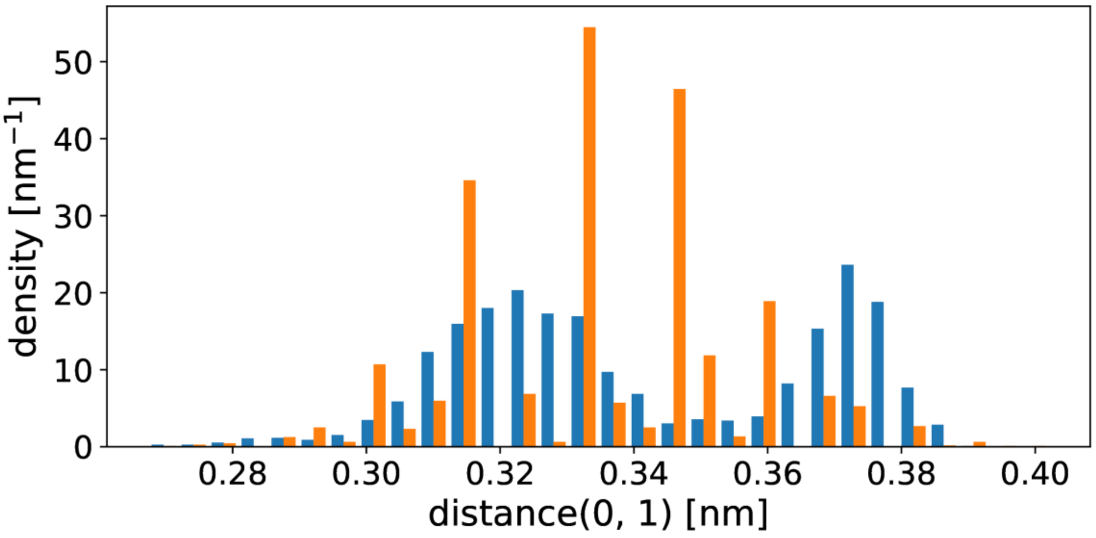


Figure S2. Probability distribution of the bead distance of GABA from the MD simulations. The blue and orange bars represent the all-atom and coarse-grained models, respectively.

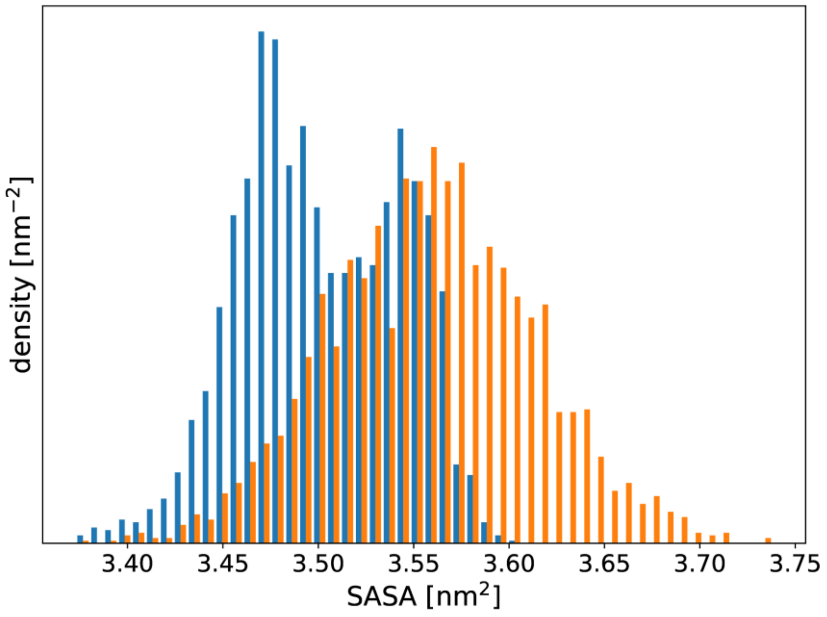


Figure S3. Probability distribution of the SASA of GABA from the MD simulations. The blue and orange bars represent the all-atom and coarse-grained models, respectively.

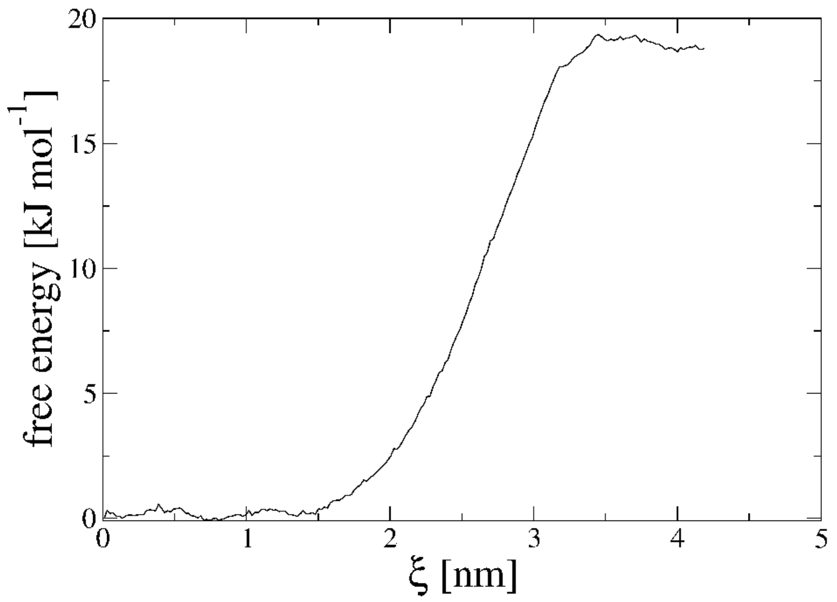


Figure S4. Free energy profile along the reaction path in the system of GABA, octanol, and water.

The results of the coarse-grained MD simulation using the propofol parameters are shown in Figs. S5-S7. The distribution of the distances between the beads reproduced the results of the all-atom model. The error of the SASA distribution was about 0.3 $nm^2$, which is sufficiently accurate. The log$P$ value calculated from the free energy profile was 3.60. This value is close to the experimental value of 3.79, indicating that the coarse-grained model reproduces the polarity of propofol.

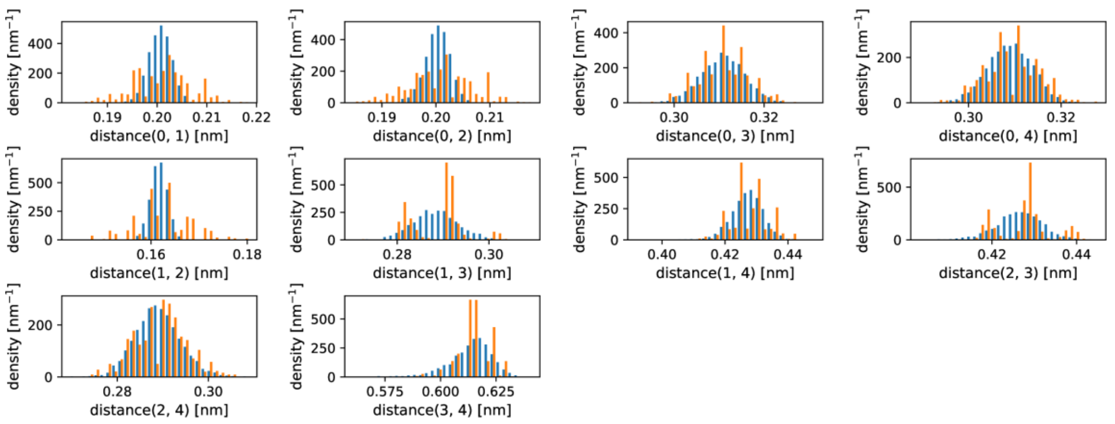


Figure S5. Probability distribution of the bead distances of propofol from the MD simulations. The blue and orange bars represent the all-atom and coarse-grained models, respectively.

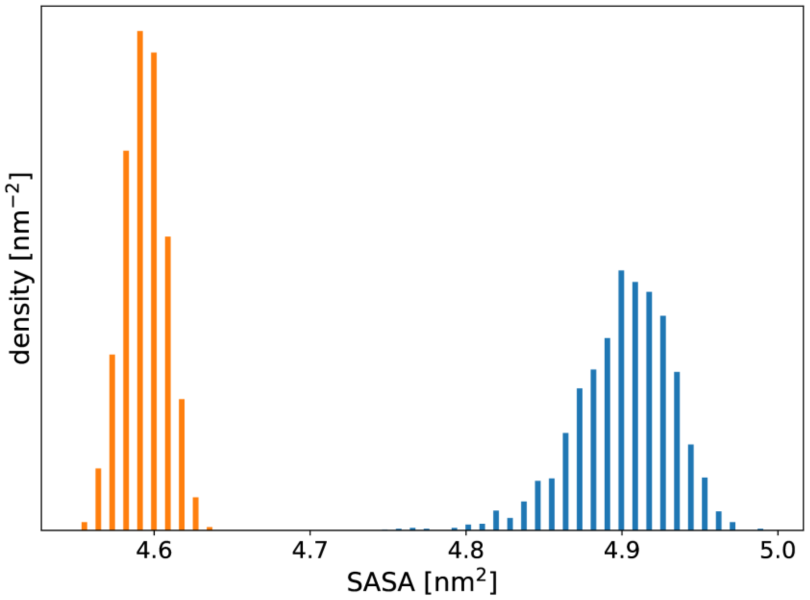


Figure S6. Probability distribution of the SASA of propofol from the MD simulations. The blue and orange bars represent the all-atom and coarse-grained models, respectively.

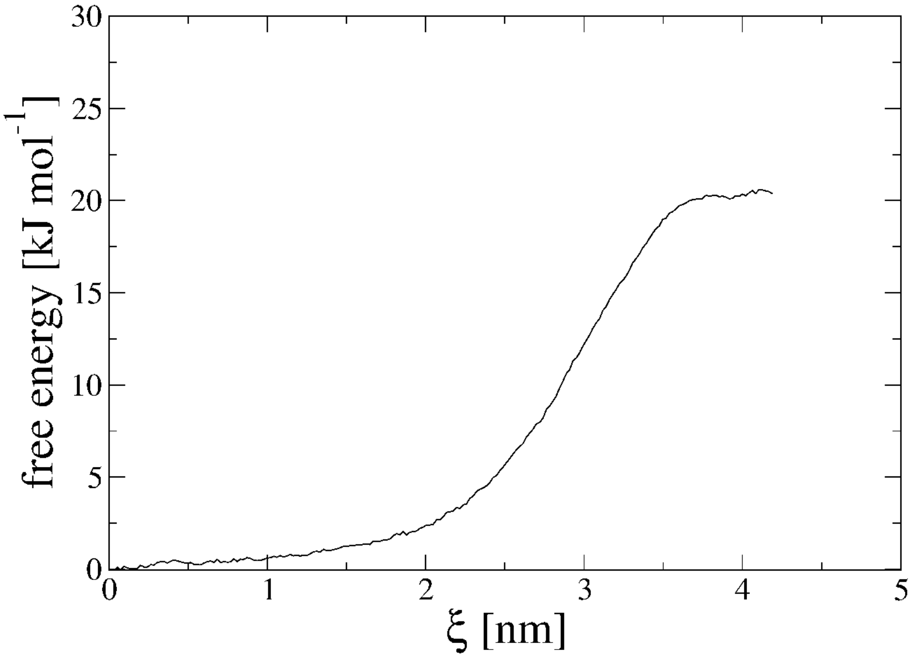


Figure S7. Free energy profile along the reaction path in the system of propofol, octanol, and water.

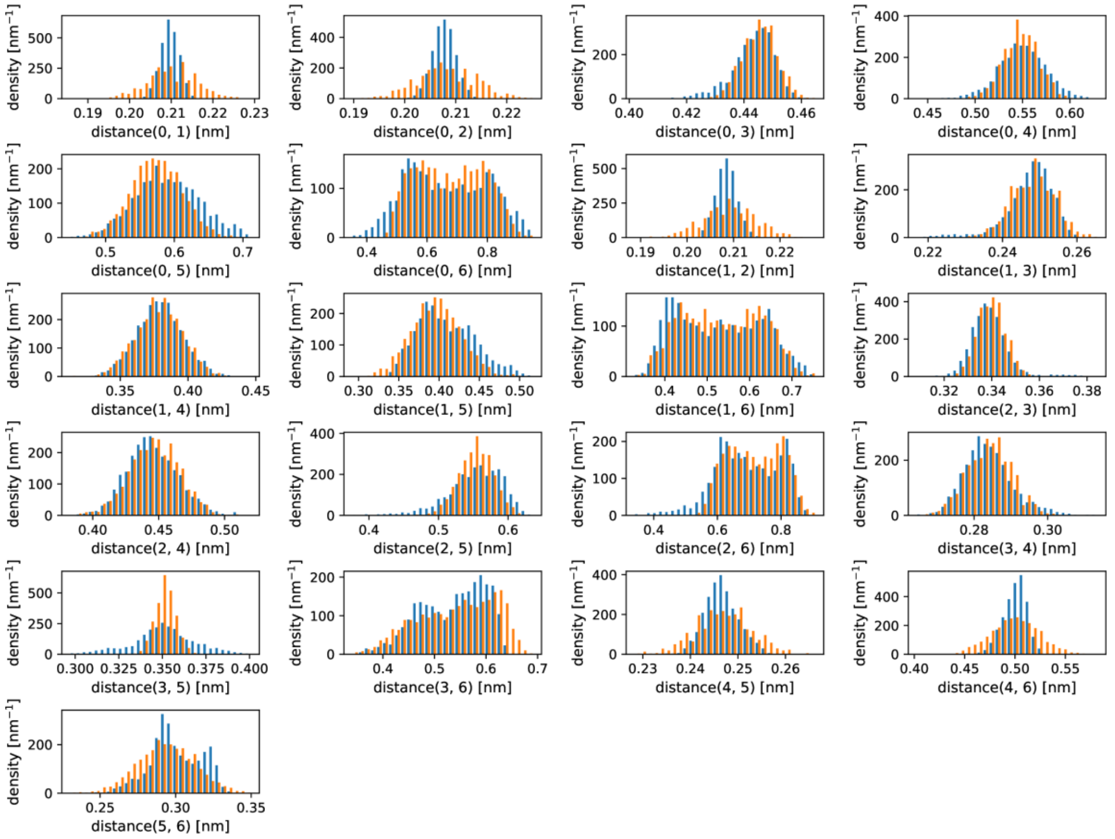


Figure S8. Probability distribution of the bead distances of etomidate from the MD simulations. The blue and orange bars represent the all-atom and coarse-grained models, respectively.

The results of the coarse-grained MD simulation using the etomidate parameters are shown in Figs. S8-S10. The distributions for the distances between the beads and SASA in the coarse-grained model reproduced the results of the all-atom model. In addition, the log*P* value calculated from the free energy profile was 3.12. This value is close to the experimental value of 3.05, indicating that the coarse-grained model reproduces the polarity of etomidate.

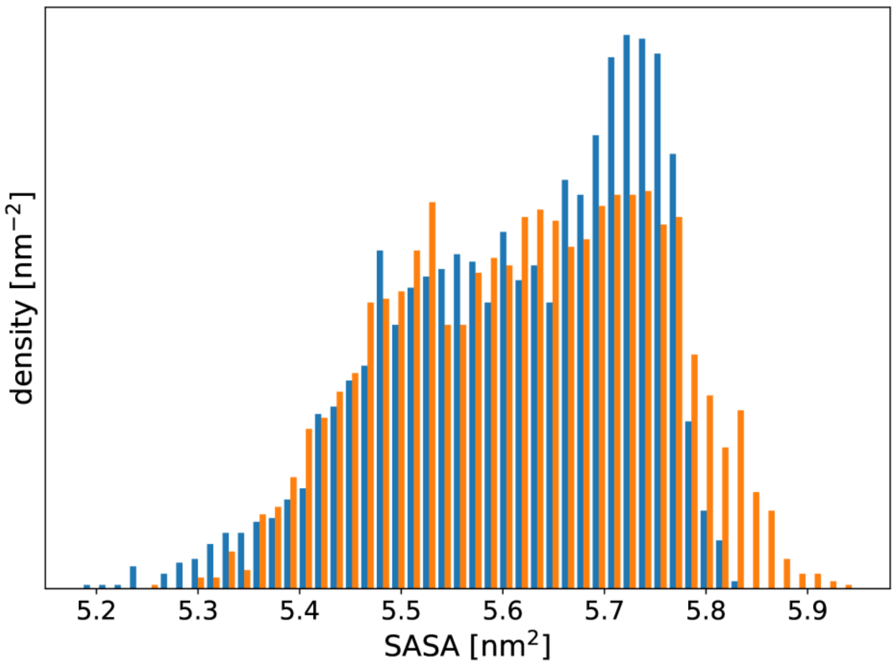


Figure S9. Probability distribution of the SASA of etomidate from the MD simulations. The blue and orange bars represent the all-atom and coarse-grained models, respectively.

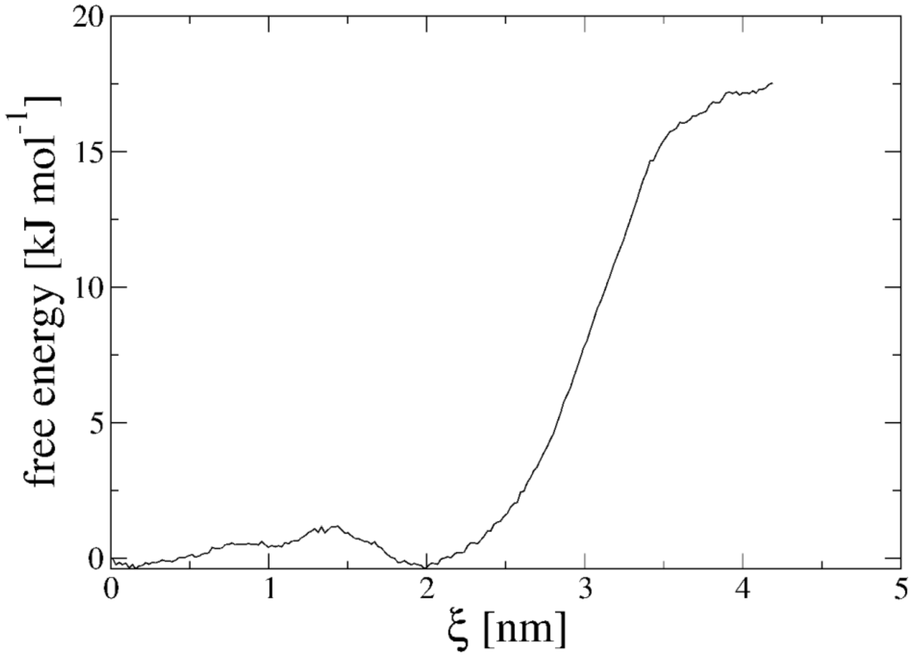


Figure S10. Free energy profile along the reaction path in the system of etomidate, octanol, and water.

### Additional results for the coarse-grained molecular dynamics simulations

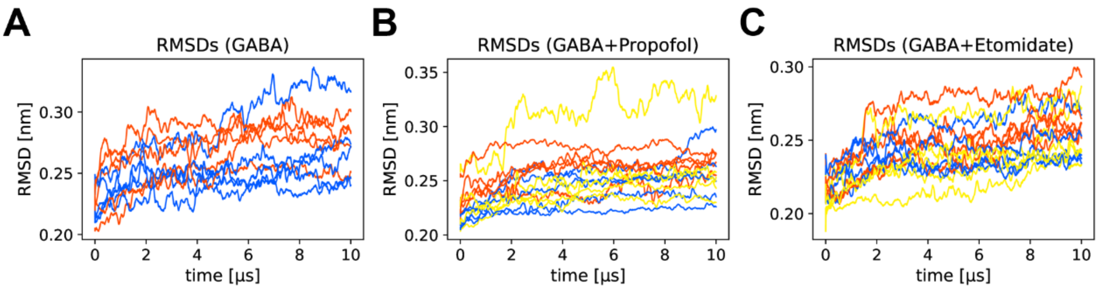


Figure S11. Time series of the RMSDs for (A) System 1, 2, (B) System 3-5, and (C) System 6-8. The yellow, orange, and blue curves correspond to the results of System 3 and 6 ($GABA_AR$ with GABA and propofol or GABA and etomidate), System 1, 4, and 7 ($GABA_AR$ with GABA), and System 2, 5, and 8 ($GABA_AR$ without ligands), respectively.

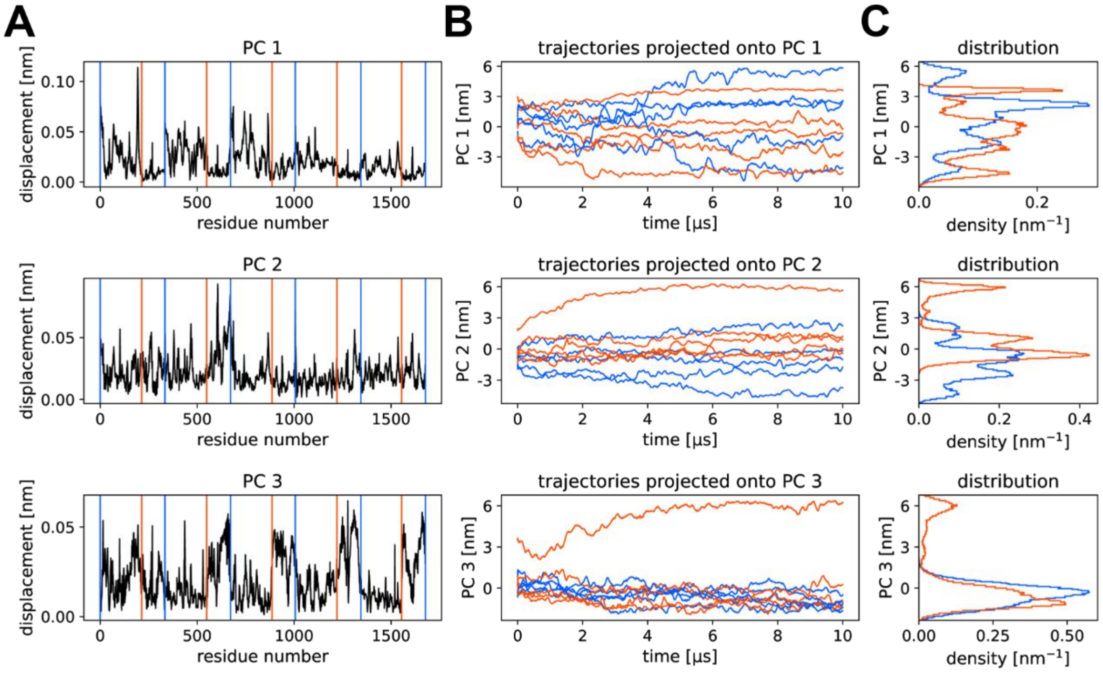


Figure S12. Results of the PCA for System 1, 2 ($GABA_AR$ with GABA). (A) The displacement of each residue is plotted for the first three PCs (The explained variance ratio of PC1, PC2, and PC3 is 0.172, 0.131, and 0.084, respectively). The blue and red vertical lines represent the subunit and ECD-TMD boundaries, respectively. (B, C) The trajectories projected onto each PC and their distribution are shown. The trajectories from System 1 (red, $GABA_AR$ with GABA) and System 2 (blue, $GABA_AR$ without GABA) were projected onto the PCs calculated from the trajectories of System 1, 2.

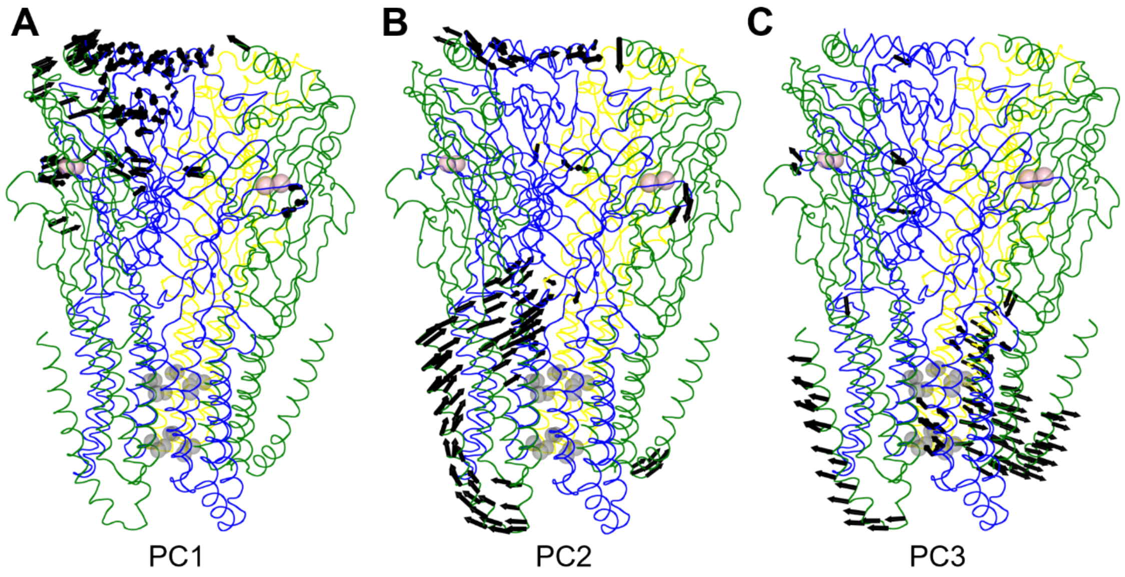


Figure S13. (A) PC1, (B) PC2, and (C) PC3 for $GABA_AR$ with GABA. The largest 100 components of each PC are shown by black arrows. The gray and pink spheres show the gate residues of the pore and GABA, respectively. The α, β, and γ subunits are colored green, blue, and yellow, respectively.

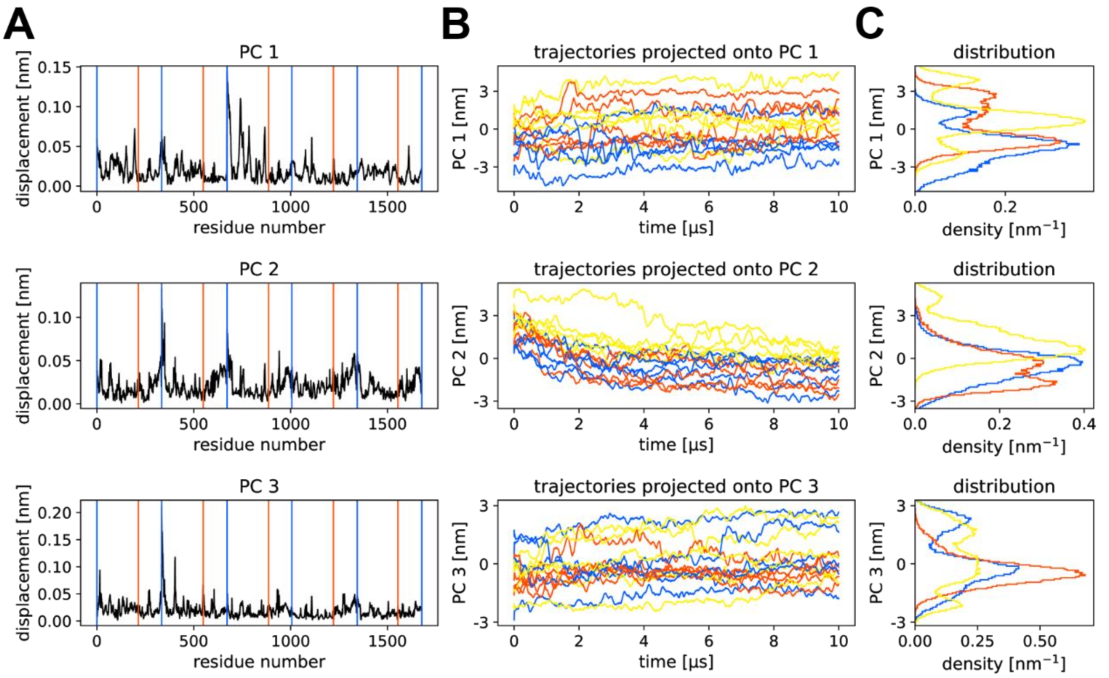


Figure S14. Results of the PCA for System 6-8 (GABA$_A$R with GABA and etomidate). (A) The displacement of each residue is plotted for the first three PCs (The explained variance ratio of PC1, PC2, and PC3 is 0.118, 0.073, and 0.055, respectively). The blue and red vertical lines represent the subunit and ECD-TMD boundaries, respectively. (B, C) The trajectories projected onto each PC and their distribution are shown. The trajectories obtained from System 6 (yellow, GABA$_A$R with GABA and etomidate), System 7 (red, GABA$_A$R with GABA), and System 8 (blue, GABA$_A$R without ligands) were projected onto the PCs calculated from the trajectories of System 6-8.

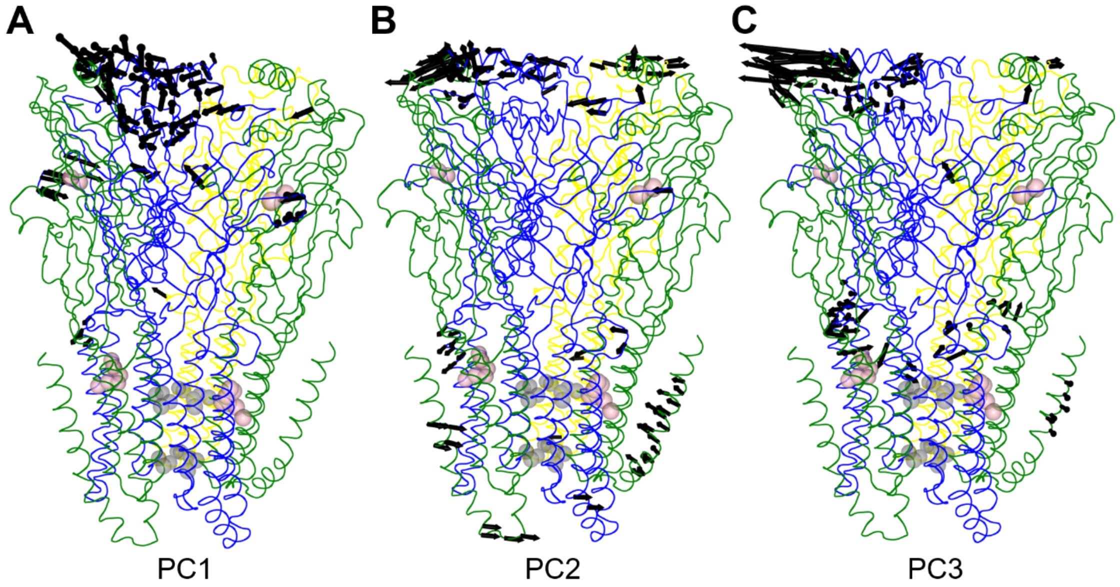


Figure S15. (A) PC1, (B) PC2, and (C) PC3 for GABA$_A$R with GABA and etomidate. The largest 100 components of each PC are shown by black arrows. The gray and pink spheres show the gate residues of the pore and the ligands, respectively. The α, β, and γ subunits are colored green, blue, and yellow, respectively.

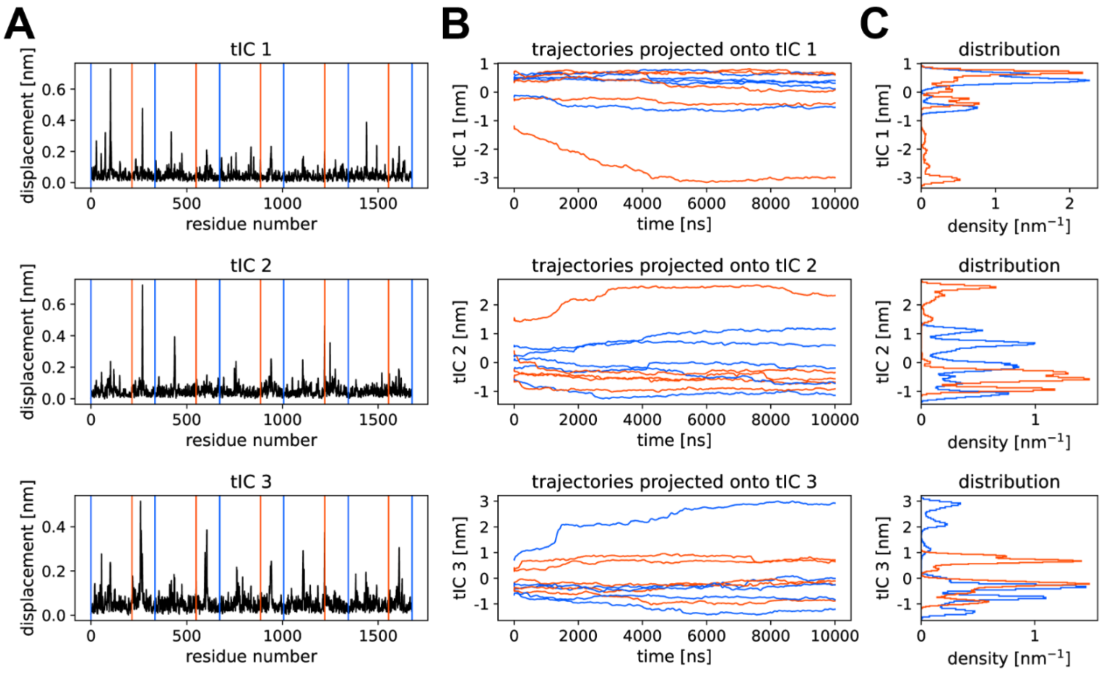


Figure S16. Results of the tICA for System 1, 2 ($GABA_AR$ with GABA). (A) The displacement of each residue is plotted for the first three tICs. The blue and red vertical lines represent the subunit and ECD-TMD boundaries, respectively. (B, C) The trajectories projected onto each tIC and their distribution are shown. The trajectories obtained from System 1 (red, $GABA_AR$ with GABA) and System 2 (blue, $GABA_AR$ without GABA) were projected onto the tICs calculated from the trajectories of System 1, 2.

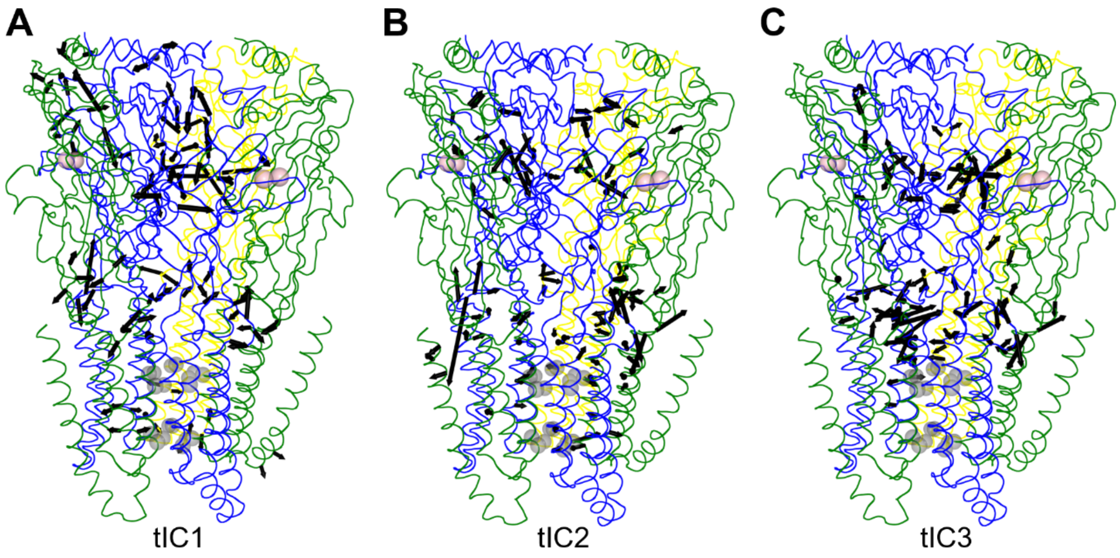


Figure S17. (A) tIC1, (B) tIC2, and (C) tIC3 for $GABA_AR$ with GABA. The largest 100 components of each tIC are shown by black arrows. The gray and pink spheres show the gate residues of the pore and GABA, respectively. The α, β, and γ subunits are colored green, blue, and yellow, respectively.

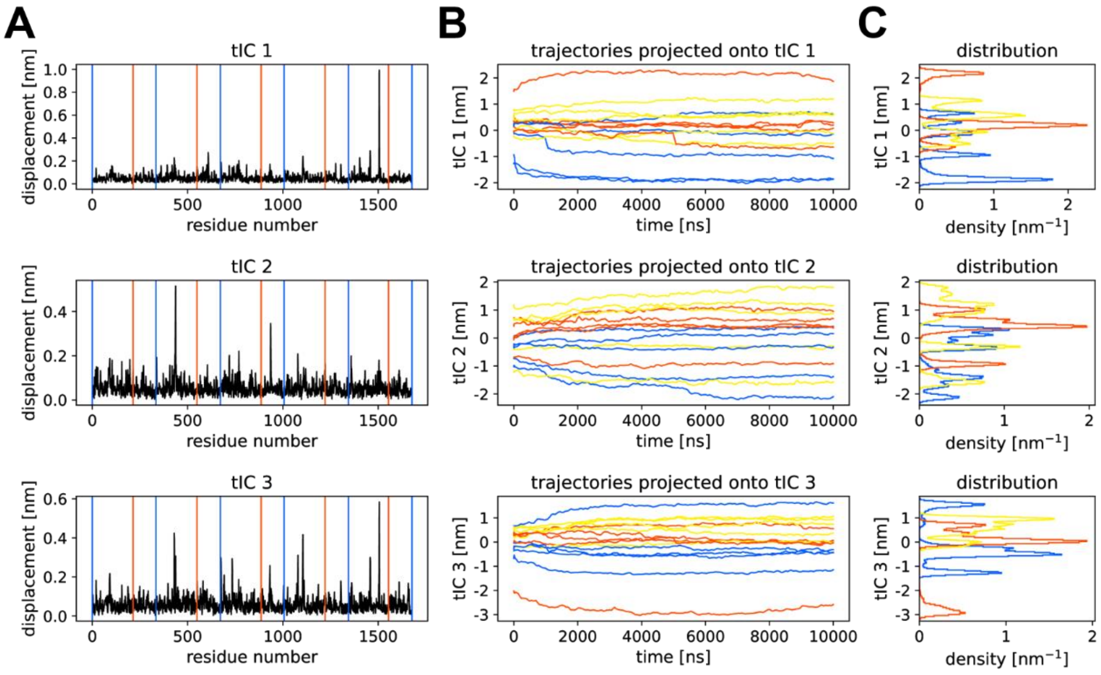


Figure S18. Results of the tICA for System 3-5 (GABA$_A$R with GABA and propofol). (A) The displacement of each residue is plotted for the first three tICs. The blue and red vertical lines represent the subunit and ECD-TMD boundaries, respectively. (B, C) The trajectories projected onto each tIC and their distribution are shown. The trajectories obtained from System 3 (yellow, GABA$_A$R with GABA and propofol), System 4 (red, GABA$_A$R with GABA), and System 5 (blue, GABA$_A$R without ligands) were projected onto the tICs calculated from the trajectories of System 3-5.

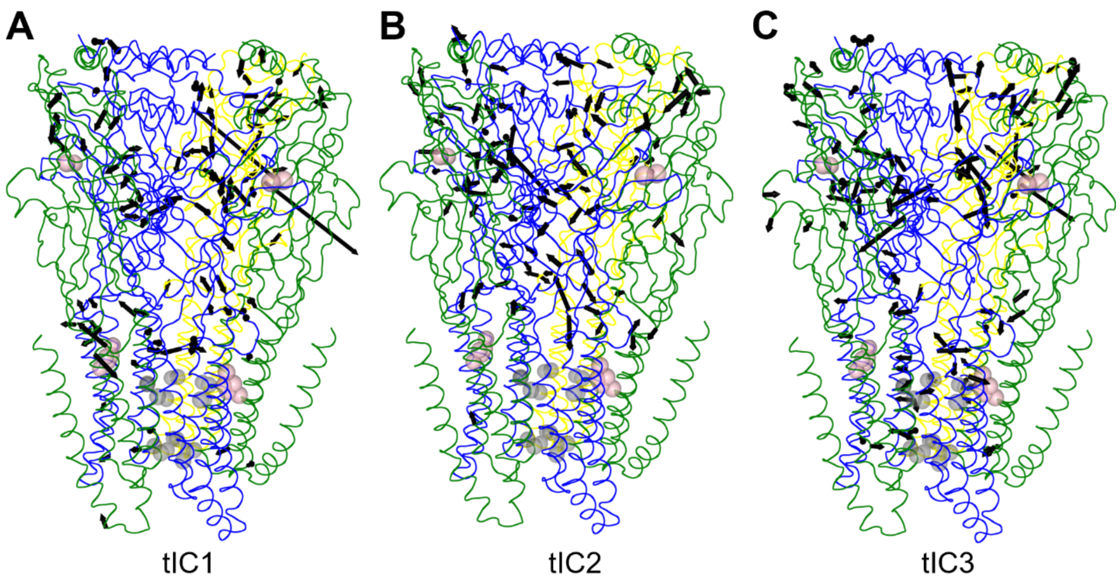


Figure S19. (A) tIC1, (B) tIC2, and (C) tIC3 for $GABA_AR$ with GABA and propofol. The largest 100 components of each tIC are shown by black arrows. The gray and pink spheres show the gate residues of the pore and the ligands, respectively. The α, β, and γ subunits are colored green, blue, and yellow, respectively.

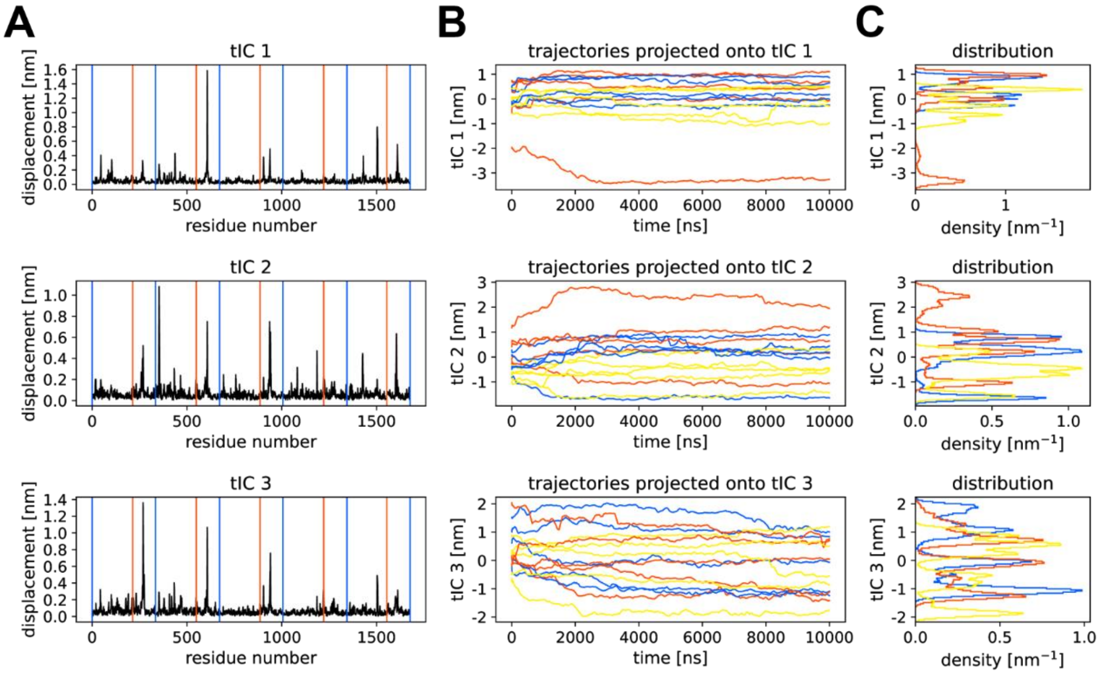


Figure S20. Results of the tICA for System 6-8 ($GABA_AR$ with GABA and etomidate). (A) The displacement of each residue is plotted for the first three tICs. The blue and red vertical lines represent the subunit and ECD-TMD boundaries, respectively. (B, C) The trajectories projected onto each tIC and their distribution are shown. The trajectories obtained from System 6 (yellow, $GABA_AR$ with GABA and etomidate), System 7 (red, $GABA_AR$ with GABA), and System 8 (blue, $GABA_AR$ without ligands) were projected onto the tICs calculated from the trajectories of System 6-8.

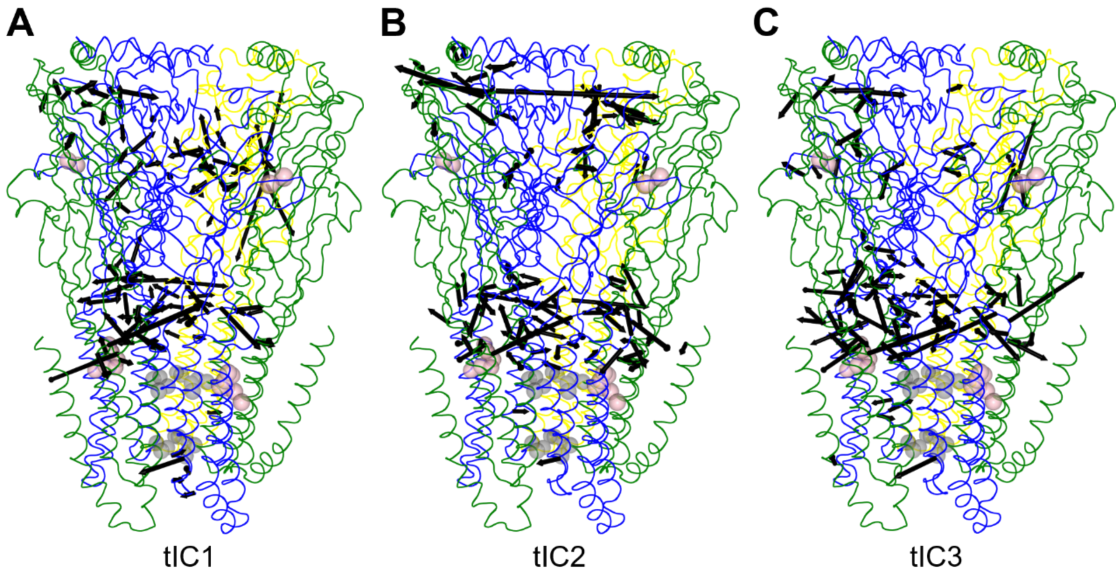


Figure S21. (A) tIC1, (B) tIC2, and (C) tIC3 for $GABA_AR$ with GABA and etomidate. The largest 100 components of each tIC are shown by black arrows. The gray and pink spheres show the gate residues of the pore and the ligands, respectively. The α, β, and γ subunits are colored green, blue, and yellow, respectively.
